# Efficient Working Memory Maintenance via High-Dimensional Rotational Dynamics

**DOI:** 10.1101/2025.09.08.674838

**Authors:** Laura Ritter, Angus Chadwick

## Abstract

Working memory (WM) is fundamental to higher-order cognition, yet the circuit mechanisms through which memoranda are maintained in neural activity after removal of sensory input remain subject to vigorous debate. Prominent theories propose that stimuli are encoded in either stable and persistent activity patterns configured through attractor mechanisms or dynamic and time-varying activity patterns brought about through functionally-feedforward network architectures. However, cortical circuits exhibit heterogeneous responses during WM tasks that are challenging to reconcile with either hypothesis. We hypothesised that these complex response dynamics could emerge from an optimally noise-robust and energetically efficient solution to WM tasks. We show that, in contrast to previous theories, networks optimised for efficient WM encoding exhibit high-dimensional rotational dynamics. We find direct evidence for these rotational dynamics in large-scale recordings from monkey prefrontal cortex. Our findings suggest that the complex and dynamic response properties of WM circuits emerge from efficient coding principles.

## Introduction

Working memory (WM) is a fundamental building block of intelligence (Baddeley, 1992). Storing items temporarily in WM enables animals to decouple motor actions from direct sensory input, facilitating flexible and adaptive behaviour and the learning of associations between temporally discontiguous events (Miller and Cohen, 2001). The neural basis of WM has long been thought to rely on persistent activity, with neurons firing both stably and selectively to specific memoranda throughout the delay between sensory input and motor response (Fuster and Alexander, 1971; Fuster, 1973; Funahashi et al., 1989; Goldman-Rakic, 1995). However, recent studies focusing on population-level neural representations during WM tasks have instead found highly dynamic neural activity patterns throughout the delay (Stokes et al., 2013; Stokes, 2015; Sreenivasan et al., 2014; Lundqvist et al., 2018; Stroud et al., 2024a; Daie et al., 2023). These findings suggest that the persistent activity hypothesis, and the mechanistic models inspired by it, are in need of revision.

Classic attractor models of WM dynamics, inspired by the persistent activity hypothesis, postulate that WM contents are stored in stable attractor states of population activity throughout the delay (Seung, 1996; Wang, 2001). In contrast, functionally-feedforward models propose that WM contents are stored in sequences of neural activity spanning the delay (Ganguli et al., 2008; Goldman, 2009; Harvey et al., 2012; Daie et al., 2023). However, neither attractor nor feedforward models can account for dynamic coding properties of WM circuits (Stroud et al., 2024a). While a hybrid model with transient feedforward amplification of external input into a stable attractor accounts for many aspects of dynamic coding (Stroud et al., 2023), attractor mechanisms are highly suboptimal in the presence of noise; recall precision deteriorates rapidly after stimulus offset due to noisy drift of the internal stimulus representation (Ganguli et al., 2008; Lim and Goldman, 2012; Burak and Fiete, 2012).

We asked whether the complex and dynamic activity patterns observed during WM tasks could be explained by a recurrent network architecture optimised for 1) robustness to neuronal noise and 2) energetic efficiency. In particular, noise introduces errors in WM tasks, and accurate WM performance therefore requires noise-resilient dynamics (Orhan and Ma, 2023; Stroud et al., 2024b; Burak and Fiete, 2012; Panichello et al., 2019). Moreover, there is considerable pressure to minimise the energic cost of neural circuit computations (Laughlin, 2001; Padamsey et al., 2022). We therefore developed a novel method to identify solutions to WM tasks that perform optimally in the presence of ongoing noise fluctuations and with minimal metabolic cost.

Surprisingly, the solutions that emerged from this method departed substantially from all previous models (Seung, 1996; Ganguli et al., 2008; Goldman, 2009; Stroud et al., 2023). In particular, we found that rotational dynamics, which are widely observed in a variety of non-WM tasks (Churchland et al., 2012; Aoi et al., 2020; Libby and Buschman, 2021), substantially outperformed the attractor and feedforward WM solutions in terms of both noise-robustness and energetic cost. These dynamics were integrated into high-dimensional feedforward-rotational motifs that closely resembled, but substantially outperformed, state-of-the-art State Space Models (SSMs) recently developed for analysis of time series data (Voelker et al., 2019; Gu et al., 2020, 2023). Moreover, these solutions recapitulated a wide range of functional properties of WM circuits, including dynamic coding at the single-neuron and population level (Spaak et al., 2017), and were present in large-scale recordings from prefrontal cortex of monkeys performing a classic spatial WM task (Panichello et al., 2024).

Our findings suggest that normative pressures on noise-robustness and energetic cost shape the complex dynamics of WM circuits. Given their similarity to SSMs, we suggest that the feedforward-rotational dynamics employed by WM circuits forms a high-dimensional substrate for flexible computation on temporally distributed data (Voelker et al., 2019; Gu et al., 2020, 2023). Beyond WM tasks, the method we develop to optimise noisy RNNs enables integration of the computation-through-dynamics framework with efficient coding theories (Averbeck et al., 2006; Sussillo, 2014; Vyas et al., 2020), allowing rigorous investigation of noise-robust and metabolically efficient dynamics across diverse cognitive tasks.

## Results

### Optimal Dynamics for Working Memory Tasks

We sought to determine the optimal dynamics for a WM task in which one of several possible input stimuli must be decoded from the response of a network after a delay, with ongoing noise inputs corrupting the stimulus representation over time (Figure 1, Figure 2A,B). For linear networks driven by Gaussian noise, the performance of an optimal decoder can be quantified as a signal-to-noise ratio (SNR, see Methods). We derived this SNR for a set of canonical linear WM models both analytically and numerically (Supplementary Figure S1, Supplementary Material). To test the optimality of these linear networks, we derived a Bayesian upper bound that holds for arbitrary nonlinear networks (Supplementary Figure S2, Supplementary Material).

**Figure 1:**
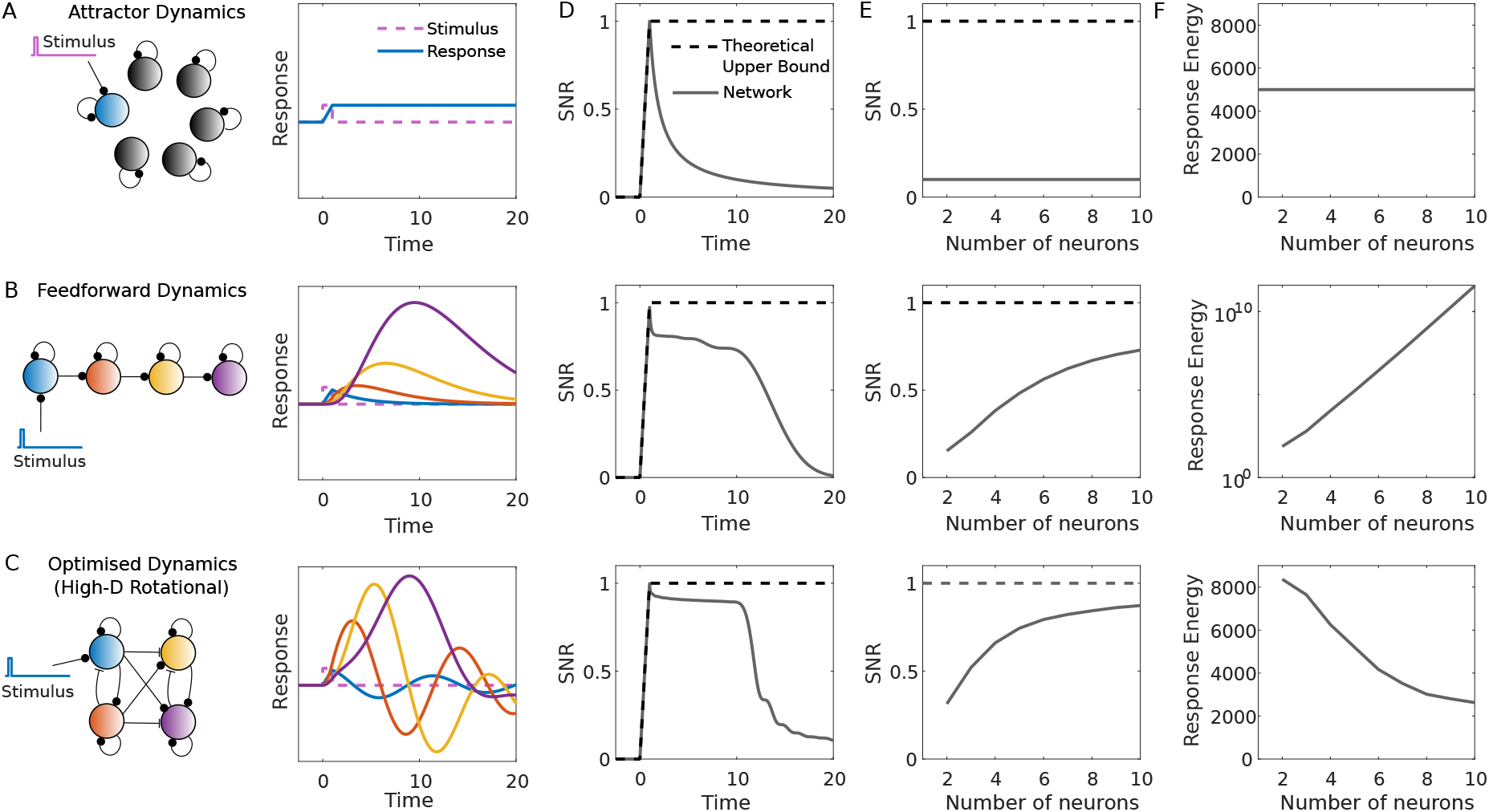
Models for working memory dynamics. A: Attractor networks generate a set of slow, non-interacting dynamical modes that store stimuli in a stable code after stimulus removal. B: Feedforward dynamics involve directed interactions over a sequence of dynamical modes, and store stimuli in a dynamic sequence of neural activity. C: Optimised networks exhibit complex, high-dimensional rotational dynamics. D: Information maintanence in each network in the presence of noise (with 10 neurons, optimised for stimulus readout at *t* = 10). Attractor networks rapidly lose information after stimulus offset due to accumulation of noise (top). Feedforward and optimised networks can achieve close to theoretically optimal decoding performance up to the optimised readout time. E: Stimulus encoding improves as the number of neurons (or dynamical modes) increases in feedforward and optimised networks, but not in attractor networks. F: The energetic cost (measured as the squared magnitude of the network response integrated over time) increases exponentially with the number of neurons for feedforward networks (note log scale on y axis), is independent of the number of neurons for attractor networks, and decreases with the number of neurons for optimised networks.

**Figure 2:**
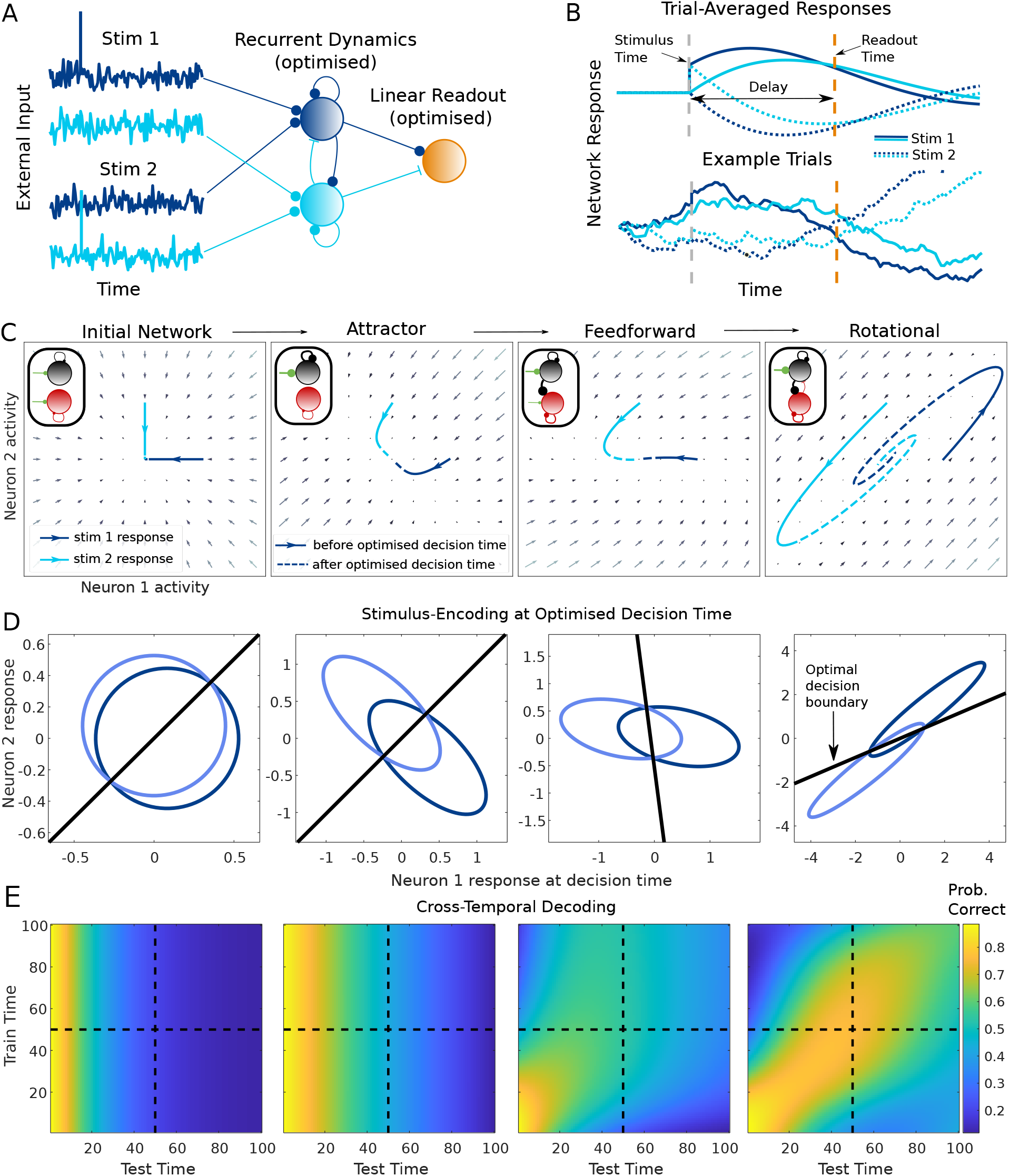
Optimisation of a two-neuron network for a WM task. A: Networks were optimised to maximise the discrimination of two noisy input stimuli after a delay. B: Responses of the optimised network to each stimulus. C: Dynamics of the network at four points during optimisation: the initial network (left), the final network before feedforward dynamics emerged (middle left), the final network before rotational dynamics emerged (middle right) and the network at convergence (right). Trajectories show the mean network response for each stimulus, with solid lines up to the optimised decision time and dashed lines thereafter. Insets show the decomposition of the network into dynamical modes (Schur decomposition). D: Response distributions under each stimulus at the decision time. E: Performance of a decoder optimised for time *t* and tested on time *t*^*′*^. Dashed lines show the optimised decision time.

We first considered the performance of two classic models for WM tasks - attractor and feedforward networks (Seung, 1996; Ganguli et al., 2008; Goldman, 2009). Attractor networks generate a set of slow, non-interacting dynamical modes that can be used to load and store information over the delay (Figure 1A, Supplementary Figure S1A). We found analytically that attractor networks are highly susceptible to noise (Figure 1D top, Supplementary Figure S1B-D), with the signal-to-noise ratio for discrimination of different input stimuli decaying exponentially after stimulus offset (Supplementary Material). In contrast, feedforward networks utilise a directed chain of dynamical modes to store a stimulus in an activity sequence that spans the delay (Figure 1B, Supplementary Figure S1E-G) (Ganguli et al., 2008; Goldman, 2009). We found that, in contrast to attractor networks, feedforward networks maintain a near-optimal SNR for a period of time after stimulus removal (Figure 1D middle, Supplementary Figure S1I-K). Moreover, for feedforward networks, but not attractor networks, the SNR increases as the number of neurons is increased (Figure 1E top vs middle, Supplementary Figure S1J). However, feedforward networks require strong, amplifying feedforward connectivity and an exponential increase in firing rates to achieve these SNR improvements (Figure 1F top vs middle, Supplementary Figure S1F-H). Thus, attractor networks are energetically cheap but highly susceptible to noise, while feedforward networks are energetically costly but substantially more noise-robust.

We asked whether solutions exist which outperform these hand-crafted attractor and feedforward networks in terms of noise-robustness and energetic cost. To this end, we optimised the recurrent connectivity of networks to achieve maximal response SNR while imposing a penalty on firing rates, using a novel method that performs gradient descent on these quantities directly (see Methods). Surprisingly, the optimised networks departed significantly from classic attractor and feedforward networks. Instead, they exhibited complex oscillatory dynamics distributed over a range of frequencies (Figure 1C). Remarkably, these networks simultaneously outperformed feedforward networks in terms of SNR and attractor networks in terms of energetic cost. As in feedforward networks, the SNR increased with the number of neurons in the network. However, unlike attractor or feedforward networks, their energetic cost simultaneously decreased (Figure 1F bottom). Moreover, the SNR of the optimised networks approached the theoretical upper bound for any (potentially nonlinear) network, suggesting that the assumption of linear dynamics was not a limiting factor.

Taken together, we find that networks optimised for WM performance exhibit a novel solution that is both noise-robust and energetically efficient, substantially outperforming classic attractor and feedforward models. We next sought to understand the dynamical mechanisms employed by these optimised networks.

### Efficient Working Memory Encoding via Amplifying Rotational Dynamics

To gain insight into the dynamical solutions employed by the optimised networks, we considered a minimal model of the WM task comprising a network of two neurons receiving input from one of two stimuli presented as an instantaneous pulse (Figure 2A,B). The network was driven to stationary state by Gaussian noise before stimulus onset and optimised to maximise the SNR for discrimination of the two input stimuli after a delay, with or without a penalty on energetic cost (see Methods).

Three distinct phases were observed as the network performance progressively converged toward an optimised solution (Figure 2C, Supplementary Figure S3). In the first phase of optimisation, an approximate line attractor was formed (Figure 2C second panel). During the second phase there was a transition from attractor to feedforward dynamics (Figure 2C third panel). The final phase showed a transition to rotational dynamics (Figure 2C final panel). Similar dynamics emerged in networks with and without a penalty on energetic cost (Supplementary Figure S3).

We asked whether this rotational solution was a specific consequence of our choice of loss function, optimisation procedure, and task setup, or a general feature of optimal dynamics for WM tasks. To test whether these findings were sensitive to the loss function and optimisation method, we investigated the solutions learned by backpropagation-through-time (BPTT) with a squared error loss, using an analytical solution in the limit of large batch size (see Methods). Remarkably, this method produced near-identical weight trajectories to those produced by optimisation of the SNR (Supplementary Figure S4). Moreover, we showed that optimising the SNR is mathematically equivalent to optimising the probability of correct choice, suggesting that incremental learning rules based on trial-to-trial feedback should lead to similar learning trajectories (Supplementary Material). To exclude other factors that may bias the networks towards rotational solutions, we investigated the influence of 1) the range of delays the network was optimised for 2) the initial network weights 3) the initial state covariance at the time of stimulus onset 4) the length of the stimulus period relative to the delay. Rotational dynamics emerged in all settings (Supplementary Figures S4-S7), including the limiting case where the stimulus was presented continuously until the decision time, i.e. an evidence integration task (Supplementary Figure S7).

The robust emergence of rotational dynamics suggests that these solutions are more noise-resilient than attractor or feedforward mechanisms. We therefore investigated how networks integrate noisy sensory input to form an efficient output population code. We plotted the distribution of noisy network responses to each stimulus at the optimised decision time (Figure 2D). Response SNR is high if these response distributions have minimal overlap (Averbeck et al., 2006). At all stages of optimisation, recurrent integration and amplification of sensory input generated elliptical response distributions with low and high variance dimensions (Figure 2D, 2nd-4th panels). Moreover, for attractor and feedforward networks, common amplification of signal and noise caused the stimulus-coding direction to be aligned to the principal noise direction (Figure 2D, 2nd and 3rd panels). Thus, attractor and feedforward networks had strong information-limiting correlations (Moreno-Bote et al., 2014). In contrast, alignment between signal and noise was reduced in the rotational network (Figure 2D, rightmost panel). Thus, the rotational network acted to both amplify the input signal and minimise corruption of the signal by amplified noise (Supplementary Figure S8).

To determine how the optimal decoder varied over the delay in each network, we implemented a cross-temporal decoding analysis (Figure 2E). For attractor networks, information was encoded along a stable coding direction that was independent of delay time. In contrast, the feedforward and rotational networks exhibited signatures of dynamic coding (Stokes et al., 2013) - decoding performance depended on the disparity between the times the decoder was trained and tested on. Thus, rotational dynamics achieves optimally noise-robust WM performance through a dynamic population code.

### Higher-Dimensional Networks Employ Feedforward-Rotational Sequences

Having characterised the optimal WM dynamics for two-dimensional networks, we next investigated the properties of higher-dimensional networks optimised to solve the task. Classic work has shown that feedforward networks are optimal for WM tasks in discrete-time networks, and can also outperform attractor networks in continuous-time networks (Ganguli et al., 2008; Goldman, 2009; Lim and Goldman, 2012). More recently, continuous-time State Space Models (SSMs) such as the Legendre Memory Unit (LMU) have been developed, which compute an online approximation of their recent input history in terms of a set of temporal basis functions and achieve state-of-the-art performance on memory tasks (Voelker et al., 2019; Gu et al., 2020, 2023). However, the optimal solution for WM tasks has not been resolved.

Numerically optimised networks exhibited a combination of feedforward and rotational dynamics. For optimised *N-dimensi*onal networks, all eigenvalues were complex, meaning that the network contained *N/2 rotati*onal planes (Figure 3A,D, Supplementary Figure S9). Although distributed across neurons, task-relevant input targeted a single dynamical mode, which elicited a feedforward cascade of activity through these rotational planes (Figure 3A,D). The LMU was based on a fundamentally different architecture, but exhibited rotational structure similar to the optimised network (Figure 3B, Supplementary Figure S9). Both the LMU and optimised networks departed substantially from classic feedforward models (Figure 3C). However, despite the similarity of the optimised networks to the LMU, the optimised networks substantially outperformed both feedforward networks and LMUs on the task (Figure 3E).

**Figure 3:**
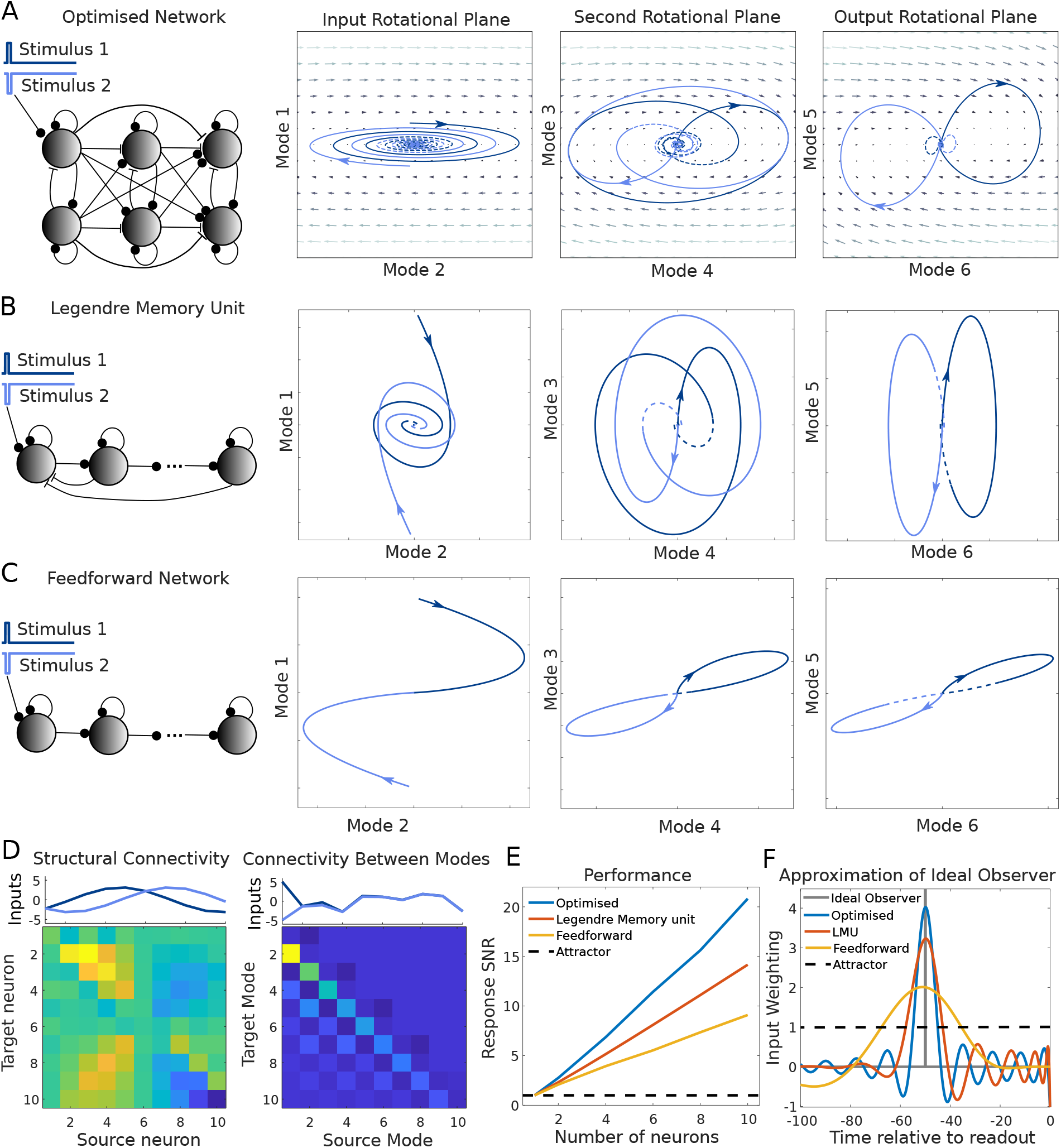
Numerically optimised higher-dimensional networks. A-C: Left: Functional connectivity between dynamical modes of an optimised 6-dimensional network (A), Legendre Memory Unit (B) and feedforward network (C). In each network, the stimulus input targets a single source mode and is cascaded through the other modes. Right: The response of each mode to two input stimuli. D: Left: The inputs (top) and weight matrix (bottom) of an optimised 10-dimensional network. Right: The Schur basis of the weight matrix, which represents interactions between the dynamical modes (as in A). E: The SNR of network output at the optimised decision time, compared to that of an optimal feedforward network, a Legendre Memory Unit and an attractor network. F: Comparison of the filter applied by the network to task inputs vs that of the Bayesian ideal observer solution. The ideal observer applies a delay filter which assigns a non-zero weight only to inputs at the time of stimulus presentation while filtering out pre- and post-stimulus noise input. The LMU and and optimised networks approximate this filter using a set of oscillatory basis functions. Attractor networks can only implement constant or exponentially decaying filters.

To understand the relative performance of these networks, we compared them to the theoretically optimal ideal observer solution (Supplementary Figure S2). This optimal solution requires the network-readout cascade to implement a delay filter, assigning a positive weight to inputs at the time of stimulus presentation and zero weight to inputs at all other times to avoid integration of pre- and post-stimulus noise. We found that only the LMU and optimised networks could successfully approximate this optimal filter, with attractor networks weighting all times equally and feedforward networks assigning positive weights to a broad range of times before and after stimulus presentation (Figure 3F). Moreover, the optimised network better approximated this ideal observer solution than the LMU, with a more sharply peaked filter around the time of stimulus presentation that enabled reduced weighting of input noise onto the readout. This sharply peaked filter was formed as a weighted combination of oscillatory modes, which provide a more flexible basis for approximating complex input-output functions than purely decaying modes.

These findings suggest that a novel dynamical mechanism combining feedforward and rotational motifs enables optimally noise-robust WM maintenance. In particular, the optimised networks compute an online approximation of their recent input history in terms of a class of oscillatory basis functions, enabling accurate approximation of a wide range of input-output functions by varying the readout weights (Voelker et al., 2019; Gu et al., 2020, 2023).

### Multiple Stimuli are Encoded in Orthogonal Rotational Subspaces

So far we have considered the optimal linear dynamics for a two-stimulus WM task (Voitov and Mrsic- Flogel, 2022; Daie et al., 2023). We next investigated WM tasks with arbitrary numbers of stimuli. As in classic delayed match to sample WM tasks, we considered a circular variable *θ ∈* [0, *2π), for* example the spatial location of a stimulus in the visual field or the colour of stimulus (Goldman-Rakic, 1995; Panichello et al., 2019). We optimised a minimal four-neuron network with cosine-tuned stimulus input (Figure 4A) for maximal response SNR after the delay.

**Figure 4:**
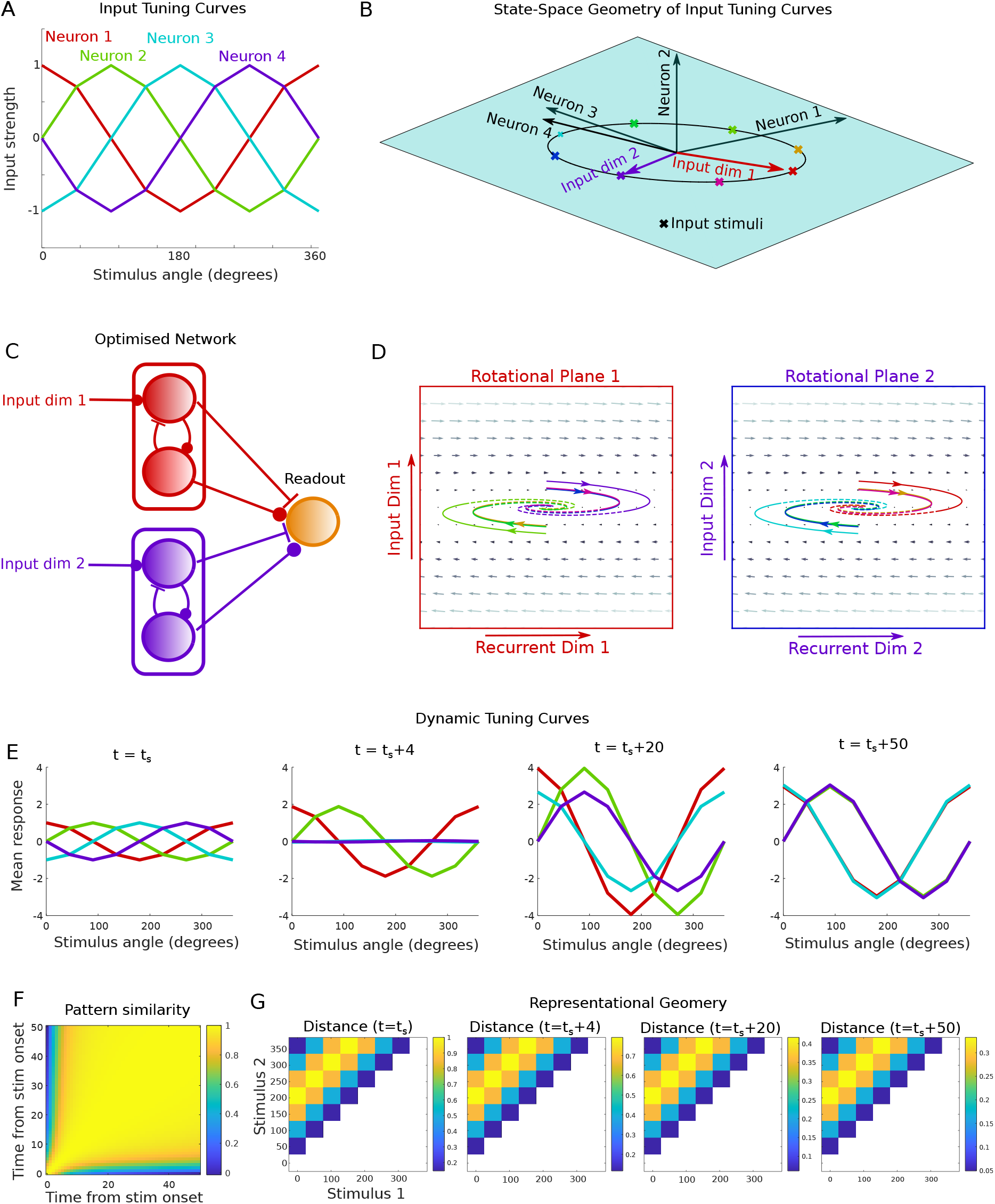
Working memory for multiple stimuli. A: Tuning curves of feedforward input to 4 neurons for a set of 8 stimuli. B: The inputs are constrained to a circle within a two-dimensional plane spanned by two basis vectors. C: The optimised network comprised two orthogonal, non-interacting subspaces, each of which integrated inputs along separate basis vector. D: Each plane exhibited rotational dynamics identical to that of networks optimised to discriminate two stimuli. Note that overlapping traces were shifted slightly for visualisation purposes. E: The response tuning curves changed dynamically over the delay, with two neurons switching to their anti-preferred stimulus. F: Cross-temporal pattern similarity analysis revealed dynamic coding early in the delay and stable coding throughout the rest of the delay. G: The representional geometry of the population response remained stable throughout the delay.

Both the stimulus inputs and network responses could be viewed within a four-dimensional state space, with the stimulus inputs constrained to a two-dimensional subspace (Figure 4B). Thus, each stimulus input could be described as a weighted sum of two orthogonal basis vectors (Figure 4B, coloured arrows). The optimised network comprised two orthogonal, non-interacting rotational subspaces (Figure 4C). Each subspace selected the input projection onto one of the two basis vectors and fed this into a private recurrent dimension via an amplifying rotational dynamics identical to that of networks optimised for two-stimulus discrimination (Figure 4D). Thus, each of the two orthogonal subspaces selectively maintained information about the magnitude of input along one of the two input dimensions, and a linear readout could reconstruct the stimulus by linearly weighting the responses within the two subspaces (see Supplementary Material).

Remarkably, this simple four-dimensional network exhibited a range of properties observed in prefrontal cortex and other brain areas during WM tasks. For example, a subset of neurons exhibited “switching selectivity”, whereby their tuning preference dynamically changed over the delay period towards their anti-preferred stimulus (Figure 4E) (Spaak et al., 2017; Libby and Buschman, 2021; Cavanagh et al., 2018). Moreover, cross-temporal pattern similarity analysis revealed dynamic coding early in the delay but stable coding throughout the remainder of the delay (Figure 4F) (Spaak et al., 2017; Stroud et al., 2024a). Finally, this dynamic code unfolded within a stable representational geometry, as estimated by the pairwise distances between population responses to different stimuli (Figure 4G)(Spaak et al., 2017).

Thus, a simple four-dimensional network optimised to perform a multi-stimulus WM task replicates wide range of properties of WM circuits via rotational dynamics in a set of orthogonal, non-interacting subspaces.

### Prefrontal Cortex Exhibits Multi-Dimensional Rotational Dynamics in a Working Memory Task

The above findings suggest that rotational dynamics is optimal for WM tasks and accounts for features of dynamic coding found in the prefrontal cortex (PFC) and other brain regions. We next asked whether signatures of rotational dynamics could be found directly in large-scale recordings of neural circuit activity during WM tasks. To this end, we analysed neuropixel recordings from PFC of monkeys performing a classic spatial WM task (Panichello et al., 2024). On each trial of the experiment, a brief (50 ms) visual stimulus was presented at one of eight spatial locations. After a delay of 1.4-1.6 seconds, the monkeys reported the remembered stimulus via an eye movement to the corresponding location.

To uncover task-related neural dynamics in the data, we implemented a novel variant of Targeted Dimensionality Reduction (TDR) (Mante et al., 2013; Aoi et al., 2020). Specifically, we modelled activity of neuron *i* at time *t* on trial *k* as

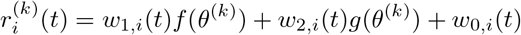

where *θ*^(*k*)^ was the stimulus presented on trial *k*. The stimulus-dependent factors *f*(*θ*), *g*(*θ*) and temporal factors *w*_*p,i*_(*t*) were fitted directly to the observed neural responses using least squares regression (Methods). The resulting model offered a compact description of the neural population responses in terms of a set of stimulus-dependent factors shared across neurons and temporal factors shared across stimuli.

To quantify the quality of model fit, we compared the post-stimulus time histograms (PSTHs) of each neuron under the TDR model to their true PSTHs. Despite being heavily underparameterised (37.5% as many TDR parameters as PSTH data points), the TDR model accurately fit the diverse PSTHs of neurons in the data (*R*^2^ = 0.73 *±* 0.06, mean*±*std over 25 recording sessions, Figure 5A). Moreover, using cross-validation to predict PSTHs on held out data, we found that a TDR model with two stimulus-dependent subspaces (*f*(*θ*) and *g*(*θ*)) was optimal (Figure 5B red bars), and performed substantially better than a baseline model in which the PSTHs formed on the training data were used to predict the PSTHs on the validation data (Figure 5B, grey bar). This suggested that the structure of the TDR model successfully captured shared stimulus- and time-dependent population structure in the data.

**Figure 5:**
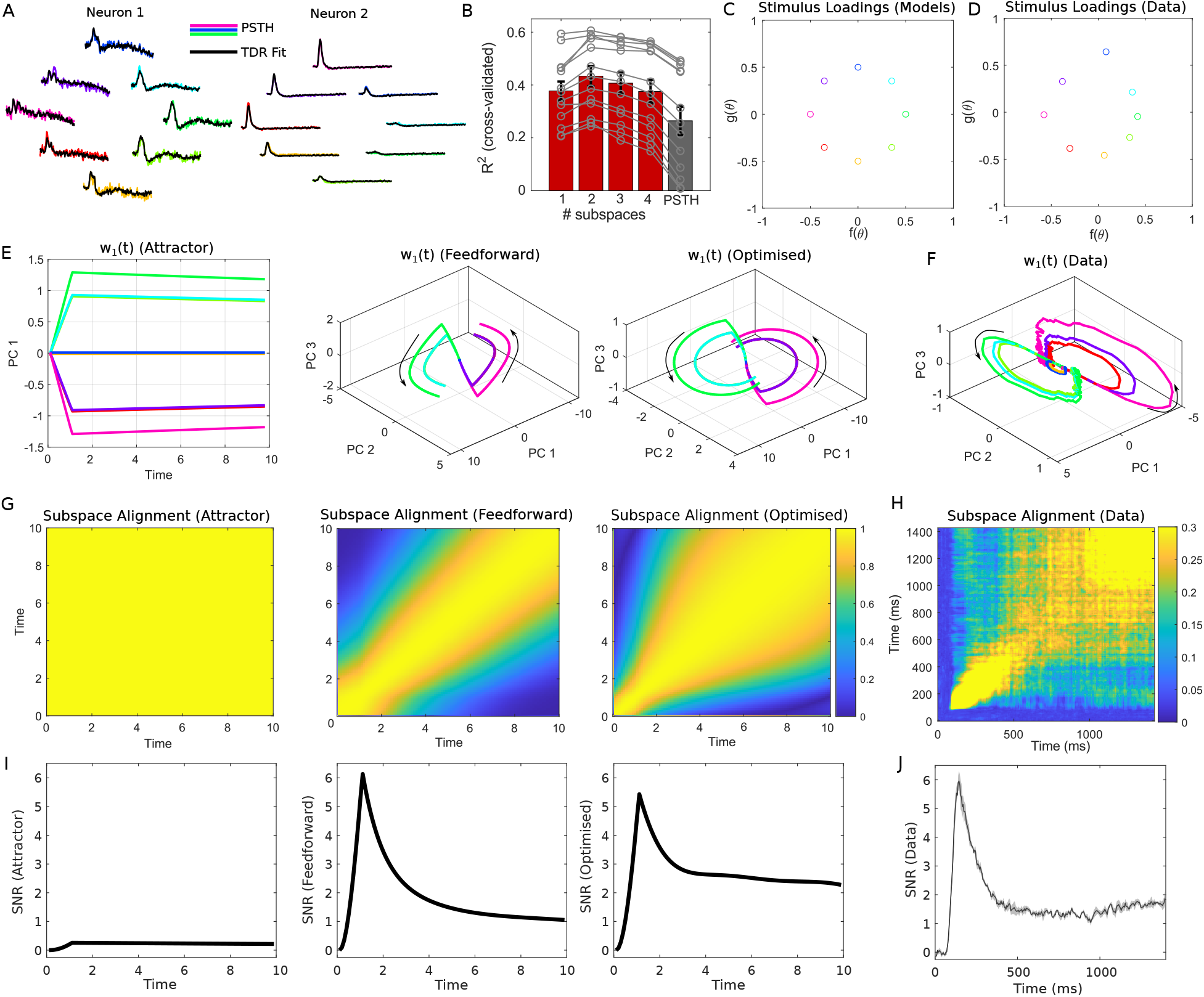
Rotational dynamics in monkey PFC during a spatial WM task. A: PSTHs of two neurons for each of the 8 stimuli in the task (coloured lines). The TDR fit is shown in black. B: Cross- validation performance of the TDR model with different numbers of stimulus components (red bars). A baseline model in which the PSTH formed on the training data is used to predict the PSTH on the test data is shown for comparison (grey bar). Error bars show mean *±* SEM over 12 sessions (sessions were excluded if the *R*^2^ for the baseline model was negative). Grey traces show individual sessions (note that the optimal number of subspaces was 2 for all included sessions). C: The stimulus factors *f*(*θ*), *g*(*θ*) learned from a TDR fit to simulated data under the models. D: The stimulus-loadings of a TDR fit to experimental data (a single recording session with 586 neurons). E: The population PSTHs (principal components of **w**_1_(*t*)) for each of the models. Note that the population PSTHs of **w**_2_(*t*) were identical to those of **w**_1_(*t*) for all models. F. As in E, but for experimental data. The second subspace **w**_2_(*t*) is shown in Supplementary Figure S10C. G: The alignment between the subspaces spanned by **w**_1_(*t*), **w**_2_(*t*) and **w**_1_(*t*^*′*^), **w**_2_(*t*^*′*^) for each of the models. H: Left: As in G, but for experimental data. I: The SNR of responses of the three models (computed on antipodal stimulus pairs). SNRs were computed assuming stationary state covariance, in contrast to Figure 1 which assumed a zero variance initial state. J: The cross-validated SNR of experimentally-recorded responses projected onto the TDR Principal Component subspace (mean*±*SEM over 4 antipodal stimulus pairs).

Given that the TDR model provided an excellent fit to the data, we next sought to compare the population structure it identified to that expected under competing models for WM dynamics. We considered classic attractor models (Seung, 1996), feedforward models (Ganguli et al., 2008; Goldman, 2009), and task-optimised models with rotational dynamics (Figures 1-4). A TDR model with two stimulus subspaces and no stimulus-independent subspace (*w*_0,*i*_(*t*) = 0) yielded an exact fit to the PSTHs generated by each WM model (*R*^2^ = 1). The stimulus factors *f*(*θ*), *g*(*θ*) were identical across all WM models considered, reflecting the circular structure of the stimulus set used in the task (Figure 5C). The temporal factors *w*_*p,i*_(*t*) evolved in low-dimensional subspaces captured by a small number of principal components (PCs) (Supplementary Figure S10B left). TDR applied to neural data revealed a similar low-dimensional encoding (Figure 5D, Supplementary Figure S10B right), albeit with an additional stimulus-independent subspace *w*_0,*i*_(*t*) not present in classic WM models (Machens et al., 2010; Cueva et al., 2020).

To investigate the structure of population activity in the low-dimensional stimulus-dependent subspaces, we plotted the top PCs of the temporal factors *w*_*p,i*_(*t*), which can be understood as low-dimensional “population PSTHs” (Figure 5E). The neural data exhibited highly rotational trajectories, which closely matched those of the rotational model and departed substantially from the attractor and feedforward models (Figure 5F). To quantify dynamic coding across models and data, we computed the cross-temporal alignment between the planes spanned by **w**_1_(*t*), **w**_2_(*t*) and **w**_1_(*t*^*′*^), **w**_2_(*t*^*′*^) (Figure 5G,H, Supplementary Figure S10D,E). When applied to the optimised rotational model and experimental data, this revealed classic signatures of dynamic coding, with the coding subspace at stimulus time and early delay period being orthogonal to that of the late delay. Attractor and feedforward models failed to replicate this dynamic code. While a recently proposed hybrid feedforward-attractor model accounted for the cross-temporal subspace alignment (Stroud et al., 2023), it exhibited lower-dimensional dynamics and population PSTHs that did not match those of the data (Supplementary Figure S10B-E).

Finally, we computed the response SNR for discrimination of antipodal stimulus pairs at each time point (Figure 5I,J). When computing the SNR we assumed that that the networks were driven to stationary state by noise input before stimulus onset. As a consequence, the SNR of an attractor network was low at all time points due to excessive integration of pre-stimulus noise along the slow attractor mode. In constrast, feedforward and optimised networks achieved substantially higher SNR, with an initial decrease after stimulus onset followed by a stable plateau (Figure 5I). Computing the cross-validated SNR of experimentally measured responses, projected into the six-dimensional TDR PC subspace, revealed a remarkably similar time course to that of the optimised model (Figure 5J).

Taken together, only the optimised model could account for the population PSTHs, dynamic coding properties, and SNR dynamics of the experimental data. Thus, neural activity in PFC during WM tasks is inconsistent with previous models involving attractor or feedforward dynamics. In contrast, rotational dynamics, emerging from directly optimised models, accurately accounts for the joint stimulus- and time-dependent neural population structure observed of PFC circuits during WM tasks.

## Discussion

We developed a novel method for optimisation of continuous-time RNNs driven by noisy inputs to solve cognitive tasks and leveraged this framework to find the optimally noise-robust and energetically efficient dynamics for working memory (WM). The solutions we identified combine functionally-feedforward and rotational dynamics, and substantially outperform all previously proposed linear models of WM (Seung, 1996; Ganguli et al., 2008; Stroud et al., 2023; Voelker et al., 2019). Moreover, these solutions provide a parsimonious explanation of a wide range of functional properties of WM circuits, including the widely observed phenomenon of dynamic coding (Stokes et al., 2013; Stokes, 2015; Spaak et al., 2017; Stroud et al., 2024a; Spaak et al., 2017; Cavanagh et al., 2018; Libby and Buschman, 2021). Thus, we suggest that WM circuits are adapted for efficient computation.

Although our theoretical analysis was restricted to linear networks, several lines of argument suggest that our conclusions apply to nonlinear networks. First, the nonlinear dynamics of PFC during simple tasks has been shown to be well approximated by a linear dynamical system (Machens et al., 2010; Soldado-Magraner et al., 2024), and the dynamical solutions of task-trained nonlinear RNNs in WM and other tasks can often be qualitatively captured by linear approximations (Mante et al., 2013; Stroud et al., 2023). Second, we derived a non-dynamical Bayesian ideal observer analysis that places strong constraints on the optimal dynamics of any (linear or nonlinear) dynamical solution to the WM task, and showed that optimal linear networks achieve this upper bound. Third, we analysed large-scale recordings from monkey PFC during a WM task using a non-dynamical TDR analysis, and showed that the population structure was consistent with optimal rotational dynamics but not classic attractor or feedforward models (Seung, 1996; Ganguli et al., 2008; Goldman, 2009) or recently proposed hybrid feedforward-attractor models (Stroud et al., 2023).

Although rotational dynamics have not previously been considered for WM tasks, they have been observed across a wide range of brain areas and tasks, including in PFC in contextual decision-making tasks (Aoi et al., 2020) and in auditory cortex in implicit short-term memory tasks (Libby and Buschman, 2021). Moreover, high-dimsensional dynamics distributed across multiple brain regions have been observed in WM tasks (Voitov and Mrsic-Flogel, 2022; Daie et al., 2023). Indeed, although we analysed experimental data collected from to PFC, we speculate that the theoretically optimal rotational dynamics we identify may be distributed across a wide network of regions of the brain that participate in the task (Christophel et al., 2017; Voitov and Mrsic-Flogel, 2022).

While, to our knowledge, feedforward-rotational dynamics have not been previously considered in the neuroscience literature, their properties can be understood in light of State Space Models (SSMs) recently developed for machine learning applications. SSMs such as the Legendre Memory Unit, MAMBA and HiPPO are built upon linear dynamical systems which are mathematically optimised to compute an online approximation of their recent input history in terms of a set of orthogonal polynomials (Voelker et al., 2019; Gu et al., 2020, 2023). We found that networks optimised for WM tasks exhibit qualitatively similar properties to these SSMs, but substanitally outperform them in the WM tasks we consider. This suggests that WM circuits may compute an online approximation of their input history in terms of an orthogonal basis of oscillatory functions. Given that SSMs now achieve state-of-the-art performance on a plethora of time-series tasks, the high-dimensional rotational solutions implemented by WM circuits may be suitable for a wide range of complex tasks beyond simple delayed recall paradigms typically studied experimentally. In particular, arbitrary functions of an SSM’s recent input history can be reconstructed through simple linear readouts, providing a flexible substrate for memory-based computation (Gu et al., 2023). This use of flexible high-dimensional representations mirrors those used by PFC for complex cognitive tasks (Fusi et al., 2016), but extends them to the time domain. Thus, we suggest that PFC dynamically computes a high-dimensional, optimally compressed representation of its input history for general-purpose computation on temporally distributed data.

## Methods

### Task Setup

We considered continuous-time recurrent linear networks driven by noisy inputs:

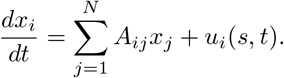

The input comprised a stimulus at time *t* = 0 and ongoing Gaussian noise *u*_*i*_(*s, t*) = ũ(*t*)*u*_*i*_(*s*) + *n*_*i*_(*t*) where 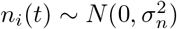. The stimulus timecourse was either a delta pulse ũ(*t*) = *δ*(*t*) (for Figures 2-4, S1-S6, S8) or a boxcar function ũ(*t*) = Θ(*t*) − Θ(*t* − *T*_cue_) where Θ(*t*) is the Heaviside step function and *T*_cue_ is the duration of stimulus presentation (for Figures 1, 5, S2, S7, S10).

For all networks, the input noise was set to *σ*_*n*_ = 1. The inputs were modelled as *u*_*i*_(*s*_*j*_) = *u*_0_ cos(*ŝ*_*i*_−*s*_*j*_), where the stimuli were *s*_*j*_ ∈ *{*0, *π/*2*}* for two-stimulus tasks and *s*_*j*_ = 2*π*(*j* − 1)*/N*_stim_ for multi-stimulus tasks, and the stimulus preferences were *ŝ*_*i*_ ∈ *{*0, *π/*2*}* or *ŝ*_*i*_ = 2*π*(*j* − 1)*/N*. We set *u*_0_ = *σ*_*n*_*/ ∥***u**(*s*_1_) − **u**(*s*_2_)∥ for two-stimulus tasks and *u*_0_ = *σ*_*n*_*/ ∥***u**(*s*_1_) − **u**(*s*_*N/*2_)∥ for multi-stimulus tasks in order to ensure that the input signal-to-noise ratio was normalised to 1. For multi-stimulus tasks, the inputs could be decomposed into basis vectors **u**(*s*) = *c*_1_(*s*)**b**_1_ + *c*_2_(*s*)**b**_2_, where **b**_1_ = cos **ŝ, b**_2_ = sin **ŝ** spanned a two-dimensional input space embedded in N-dimensional state space.

We analysed the SNR of an optimal linear readout of the stimulus, given by SNR 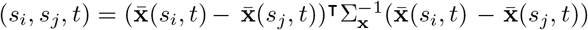 where 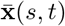 is the mean response (over trials) of the network to stimulus *s* at time *t* and Σ_**x**_ is the covariance of **x**(*s, t*) (which was independent of *s*). Note that, for the linear-Gaussian networks we considered, this linear readout is a Bayes optimal decoder of the stimulus from **x**(*s, t*) and the SNR is monotonically related to the probability of correct choice as 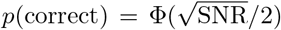, where Φ is the cumulative function of the standard normal distribution. Moreover, for continuous estimation tasks (rather than discrete classification tasks) the SNR is closely related to the Fisher information for estimation of the stimulus *s* from the response **x**. In such tasks, the SNR places a lower bound on the mean squared decoding error.

The SNR was computed directly, without sampling noisy trajectories of the system, using a Lyapunov equation to compute the covariance matrix and matrix exponentials to compute the mean response to each stimulus (see Supplementary Material). We also analysed the response energy of the network, defined as 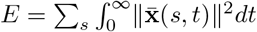, which can be computed directly in a similar manner to the SNR.

### Attractor and Feedforward Network Models

Hand-crafted networks were constructed via their Schur form *A* = *QT Q*^⊺^ where *Q* is a unitary matrix and *T* is a lower triangular matrix. For two-stimulus tasks, *Q* was constructed such that *Q*_:,1_ = **b** and all other columns of *Q* were orthogonal, with **b** = (**u**(*s*_2_) − **u**(*s*_1_))*/∥***u**(*s*_*2*_) − ***u****(s*_*1*_)∥ the input linear discriminant. For multi-stimulus tasks, *Q containe*d the two orthogonal basis vectors spanning the input space **b**_1_, **b**_2_. Specifically, we set *Q*_:,1_ = **b**_1_ and *Q*_:,*N/*2+1_ = **b**_2_, with the remaining columns orthogonal. In this case, *T* was divided into four equal-sized blocks, *T*_MultiStim_ = [*T*, 0; 0, *T*], with *T* equal to that of the two-stimulus model (described for each network below). Responses of networks were often plotted in the Schur basis **x**_Schur_(*t*) = *Q*^⊺^**x**(*t*).

For attractor networks, *T* was diagonal, with *T*_*ii*_ = *λ*_*i*_ = −1*/τ*_*i*_ where *λ*_*i*_ are the eigenvalues and *τ*_*i*_ are the decay time constants of the dynamical modes. The network had a slow dynamical mode *τ*_1_ = *τ*_slow_ and fast dynamical modes *τ*_*i>*1_ = *τ*_fast_, which with the above choice of *Q* ensured that the input linear discriminant was loaded onto the slowest dynamical mode (Chadwick et al., 2023).

Feedforward networks were constructed as *T*_*ij*_ = − *τ* ^*−*1^*δ*_*ij*_ + *ωδ*_*i,j*+1_, where *τ* is the time constant of each mode and *ω* is the feedforward weight from mode *i* − 1 to mode *i*. This ensured that the input linear discriminant was loaded onto the source mode of *T*.

The hybrid feedforward-attractor model in Supplementary Figure S10 differed from the feedforward model in two ways. First, we included only one feedforward connection. Second, this feedforward connection fed from a rapidly decaying mode into a slow attractor mode (Stroud et al., 2023). In particular, we set *T*_22_ = −1*/τ*_slow_, with *T*_*ii*_ = −1*/τ*_fast_ for *i* ≠ 2. The feedforward connection was set as *T*_21_ = *ω*.

For Figure 1, we considered a two-stimulus task. Networks had a fixed initial state **x**(*s*_*i*_, 0) = 0 (i.e., with zero initial state covariance) and a stimulus was presented for *T*_cue_ = 1 (in arbitrary units of time). The SNR for stimulus discrimination was calculated at the decision time *t*_*d*_ = 10. Attractor networks had *τ*_slow_ = 10^4^. Feedforward networks had *ω* = 5 and *τ* selected to maximise SNR.

For Figure 3, we considered a two-stimulus task in which networks were driven to stationary state by noise input before stimulus onset and received a delta pulse input, with *t*_*d*_ = 50. The SNR of attractor and feedforward networks was computed analytically using their optimal parameters (see Supplementary Material). However, because the optimal feedforward network has infinitely strong feedforward weights, we instead plotted the trajectories of this network with *τ* = 6.25, *ω* = 0.5 (where *τ* was selected as the optimal value in the limit *ω* → ∞).

For Figures 5 and S10, we considered an eight-stimulus task, with *T*_cue_ = 1 and *t*_*d*_ = 10. Attractor networks had *N* = 6 neurons, with slow time constants *τ*_slow_ = 100 and fast time constants *τ*_fast_ = 1. Feedforward networks had *N* = 12 neurons with *τ* = 5 and *ω* = 0.5. Hybrid feedforward-attractor models had *N* = 6 neurons with *τ*_fast_ = 1, *τ*_slow_ = 100 and *ω* = 1. Note that the results we present are independent of *N* for attractor and hybrid feedforward-attractor models (if *N* ≥ 2 and *N* ≥ 4 respectively), but not feedforward models.

### Optimised Networks

We minimised the following loss function:

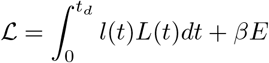

where the first term quantifies the decoder performance *L*(*t*) = Σ _*i>j*_ 1*/*SNR(*s*_*i*_, *s*_*j*_, *t*) weighted over delays by *l*(*t*) and the second term quantifies the energy of responses weighted by the regularisation parameter *β*. The weighting function was set as *l*(*t*) = *δ*(*t* − *t*_*d*_) (where *t*_*d*_ is the decision time), with the exception of Supplementary Figure S4 where we set *l*(*t*) = 1 for 0 ≤ *t ≤ t*_*d*_ and 0 otherwise. We set *β* = 10^*−*6^ for Figures 1 and S3 and *β* = 0 otherwise. For delta input tasks we set *t*_*d*_ = 50, and for finite cue tasks, we set *T*_cue_ = 1, *t*_*d*_ = 10 (with the exception of Supplementary Figure S7, where *T*_cue_ was systematically varied).

We computed the gradient 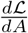 analytically and performed gradient descent 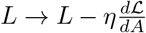 numerically with a fixed step size *η* that was set manually for each network. Network weights were initialised as *A*_*ij*_ = *δ*_*ij*_0.05(1 − 0.005*i*), except for Supplementary Figure S5 where *A* was intialised randomly using the eigendecomposition *A* = *V DV* ^*−*1^ with *V* = [cos *θ*_1_, cos *θ*_2_; sin *θ*_1_, sin *θ*_2_] with *θ*_*i*_ ∼ Unif[−*π, π*] and *D* = [*λ*_1_, 0; 0, *λ*_2_] with *λ*_*i*_ ∼ Unif[−0.2, 0], i.e. with random (real) eigenvectors and eigenvalues.

To transform the optimised networks from neuron space to the space of dynamical modes, we performed a real Schur decomposition on the weight matrix *A*_Schur_ = *Q*^⊺^*AQ*, **u**_Schur_ = *Q*^⊺^**u** where *Q* is an orthogonal matrix and *A*_Schur_ is a block upper-triangular matrix. Given that the Schur decomposition is non-unique, we exploited a set of invariances to transform the Schur decomposition into a standardised form (see Supplementary Material for details).

### Analysis of Learning Dynamics

We selected networks at four stages of optimisation to plot the flow fields, Schur modes, and responses:

1) the initial unoptimised network 2) the initial attractor network, before non-normality had increased 3) the final network before eigenvalues became complex 4) the network at convergence.

For each iteration of optimisation, we computed the SNR (defined above) and the projection of the mean and covariance of responses onto the unit length readout vector 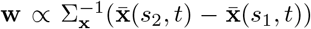, given by 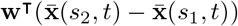 and **w**^⊺^Σ_**x**_**w** respectively (e.g., Supplementary Figure S3B,C).

The decay time constants and rotational periods of dynamical modes were computed as as *τ* = −1*/*ℛ(*λ*) and *ν* = 2*π/*ℐ(*λ*) respectively, where *λ*_*i*_ = ℛ(*λ*) *± i*ℐ(*λ*) were the eigenvalues of *A*. The alignment of modes with the input discriminant **b** were computed using either eigendecomposition or Schur decomposition. For two-neuron networks with real eigenvalues, we computed the dot product between each unit length left eigenvector of *A* (defined as *A*^⊺^**v**_*i*_ = *λ***v**_*i*_) and the input linear discriminant **b**. For higher-dimensional networks or networks with complex eigenvalues, we computed the dot product between the Schur vectors (columns of *Q*) and **b**. The non-normality of the dynamics was computed as 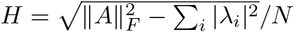, where 0 ≤ *H* ≤ 1 and *H* = 0 for a normal (attractor) network.

To optimise networks via backpropagation through time, we used analytically derived gradients in the limit of infinite batch size (Supplementary Figure S4, see Supplementary Material for derivation). We used a squared error loss ℒ = Σ_*s*_⟨∥**w** · **x**(*s, t*) + *c* − *s*∥^2^⟩ (where the expecation was taken over trials) and optimised *A* using gradient descent 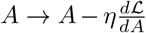 with a fixed step size *η*. The readout parameters **w**, *c* were obtained analytically at each iteration by setting 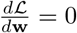 and 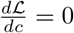.

### Analysis of Dynamic Coding

In Figure 2D, we plotted ellipses of one standard deviation of the response distribution 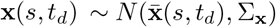 for networks at each of the four stages of optimisation we considered.

For Figure 2E, we performed a cross-temporal decoding analysis on networks at different stages of optimisation using standard signal detection theory measures. Labelling *s*_1_ = 0, *s*_2_ = 1, the optimal decoder uses a binarised linear readout *ŝ*(*c*, **w**, *t*) = Θ(**w**(*t*)^⊺^**x**(*s, t*) + *c*(*t*)), with 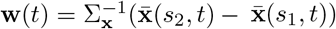 and 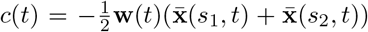. Using an arbitrary (suboptimal) **w** and *c* results in the probability of correct choice for each stimulus 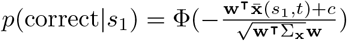 and 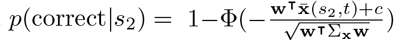 where Φ is the cumulative distribution function of the standard normal distribution. Assuming *p*(*s*_1_) = *p*(*s*_2_), the overall probability of correct choice is then 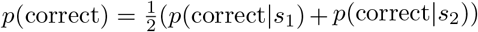. Using these formulae, we computed the probability of correct choice using the optimal decoder for time *t* on responses at time *t*^*′*^.

For Supplementary Figure S8, the signal, noise, and signal-noise-alignment were computed for networks at each stage of optimisation as signal 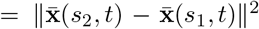, noise = Trace(Σ_**x**_), 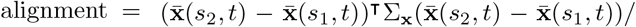 signal. The SNR was computed for optimised networks numerically (using a Lyapunov equation to compute the covariance) and the optimal SNRs of the attractor, feedforward and rotational networks were plotted using analytically-derived expressions (see Supplementary Material).

To replicate the analyses performed on experimental data by (Spaak et al., 2017), we computed the cross-temporal pattern similarity 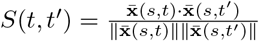 (the result was independent of the stimulus *s* for the models we considered). In addition, we computed the representational dissimilarity matrices *R*_*ij*_ = SNR(*s*_*i*_, *s*_*j*_, *t*) (Kriegeskorte and Wei, 2021).

### Legendre Memory Unit

The Legendre Memory Unit is governed by the equations

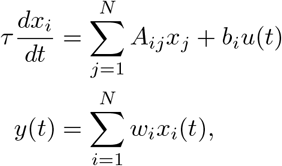

where *u*(*t*) is a scalar input time series, *τ* is a global network time constant, *A*_*ij*_ are the recurrent network weights, *b*_*i*_ is an input loading vector and *w*_*i*_ are the readout weights (Voelker et al., 2019; Voelker, 2019). Classic results from control theory provide methods to optimise the parameters of the above system such that the output *y*(*t*) is related to the input *u*(*t*) by a desired transfer function *f*, i.e. *y* ≈ (*u* ⋆ *f*)(*t*) where ⋆ is convolution (Partington, 2004; Brockett, 2015). Voelker (2019) considered the case of a delay transfer function 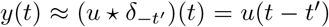, which we show in the Supplementary Material is a critical component of the Bayesian ideal observer solution to the workingmemory task we consider (Supplementary Figure S2). In approximating the delay transfer function using the above SSM, it was further shown that the solution can be understood through the lens of function approximation theory - each element of the network state *x*_*i*_(*t*) in the optimised system computes the coefficient of a Legendre polynomial approximation to the input time series *u*(*t*^*′*^) over an interval *t − τ ≤ t*^*′*^*≤ t*, and subsequent work has shown how this approach can be extended to a wide range of approximating function classes (Gu et al., 2020, 2023). As a consequence, it is possible to reconstruct arbitrary functions of the input time series over this interval through an appropriate choice of readout weights *w*_*i*_, as described below.

The parameters of the LMU are found by forming a Padé approximation of the desired transfer function in the Laplace domain (Partington, 2004; Brockett, 2015; Voelker et al., 2019). The resulting system can be written in multiple equivalent bases, all of which implement an identical transfer function. In the standard Legendre basis of the LMU, the recurrent weight matrix is given by:

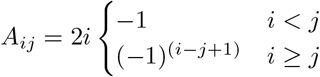

and the input vector is given by *b*_*i*_ = (2*i*)(−1)^*i−*1^. In this basis, the activations *x*_*i*_(*t*) represent the delayed inputs as 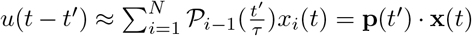 where 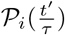 is the *i*th shifted Legendre polynomial defined on the interval 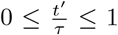 (Voelker et al., 2019). As a consequence, an arbitrary filtering of the input time series can be approximately recovered by setting the readout weights as 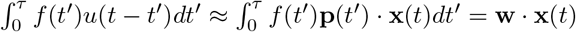 where 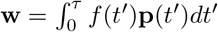.

The controllable canonical form provides an alternative representation of the LMU, where the weight matrix is given by *A*_*i*1_ = −*ν*_1_, *A*_*i,i* − 1_ = *ν*_*i*_, with *A*_*ij*_ = 0 otherwise, and the inputs are given by *b*_*i*_ = [*ν*_1_, 0, …, 0]^⊺^ (where *ν*_*i*_ = (*N* + *i −*1)(*N − i* + 1)*/i* for *i* = 1…*N*). In this basis, the inputs target only the first neuron of the network, consistent with the Schur basis for the optimised and hand-crafted networks. We therefore presented results in controllable canonical form in the main text, and compared to results in the Legendre basis in the Supplementary Material. However, we note that the two forms of the LMU are fully equivalent, with identical eigenvalues and SNR.

We simulated responses of the LMU for varying numbers of units *N* and varying time constant *τ*. Inputs comprised a delta pulse with ongoing Gaussian noise *u*(*s, t*) = *δ*(*t*)*u*(*s*) + *n*(*t*). We set *n*(*t*) ∼ *N*(0, 1) and *u*(*s*) = ±*u*_0_, with *u*_0_ chosen to normalise the input SNR to 1. We computed the mean and stationary state covariance of responses 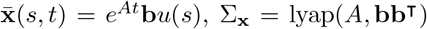 and the 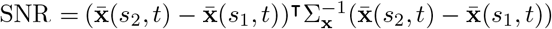. For each *N*, we chose the value of *τ* that maximised this SNR. We note that the task faced by the LMU is simpler than that faced by the other models we considered; for the LMU, input noise targets only the first unit, since **b** ∝ [1, 0, …, 0]^⊺^. Indeed, when simulating LMUs with noise targetting all units (as was the case for all other networks), performance was substantially degraded and decreased with *N* (not shown).

### Analysis of Electrophysiological Data

We analysed publicly available data from (Panichello et al., 2024). Stimuli were presented for 50 ms, followed by a delay of 1400-1600 ms after which monkeys were cued to report the remembered stimulus. We analysed data during the period from stimulus onset (0 ms) to the end of the shortest delay (1400 ms). As preprocessing steps, the raw spiking data were converted into spike counts within 25 ms bins using a sliding window of step size 5 ms. We discarded error trials and subtracted the mean spike count of each neuron (averaged over both time bins and trials). All analyses were applied to individual experimental sessions and repeated over multiple sessions to verify consistency. For Figure 5F,H,J, we used session *22-10-21*. For Supplementary Figure S10, we used sessions *22-10-24, 22-10-21* and *22-10-19*.

We fitted the targeted dimensionality reduction (TDR) model

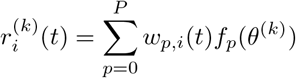

to the mean-subtracted spike counts 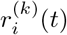 using alternating least squares, iteratively solving for *w*_*p,I*_ and *f*_*p*_ until convergence. We set *f*_0_(*θ*) = 1 for the stimulus-independent subspace. In this notation, *f*_1_(*θ*) = *f*(*θ*), *f*_2_(*θ*) = *g*(*θ*), *f*_0_(*θ*) = 1 recovers the *P* = 2 model described in the main text. Given the 8 stimuli in the experimental task, the *f*_*p*_(*θ*) parameters were given by 8-dimensional vectors *f*_*p*_ = [*f*_*p*_(*θ*_1_), …, *f*_*p*_(*θ*_8_)]^⊺^ that were inferred directly from the data. We initialised *f*_*p*_(*θ*) ∼ *N*(0, 1) (for *p >* 0) and began the optimisation algorithm by updating *w*_*p,i*_(*t*). For *N*_stim_ = 8 stimuli, *T* time points and *N* neurons in a given experimental session, the model had (*P* + 1)*NT* + *PN*_stim_ parameters, with *NT* parameters for each of the *w*_*p,i*_(*t*) components and *N*_stim_ parameters for each of the *f*_*p*_ components. In contrast, the PSTHs of the data contained *N*_stim_*NT* datapoints. This reduction in parameters reflects the shared structure of the TDR model across stimuli, which incorporates the assumption that the population PSTHs for different stimuli should have a similar temporal profile.

For cross-validation, we split the data into two subsets of randomly selected trials of equal size. The TDR model was fit to each split and used to predict the remaining split. We reported the cross-validated coefficient of determination

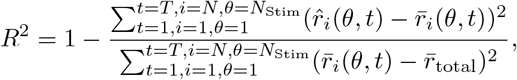

where 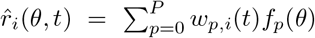 is the predicted PSTH formed by the TDR fit to the train-ing data, 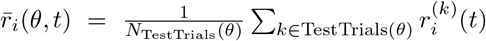 is the PSTH formed on the test data and 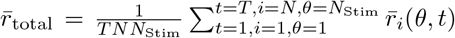. Given that the TDR model was underparameterised, we also reported this measure without cross-validation (where 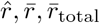 were formed on the full dataset). We repeated this cross-validation for 25 random two-way splits of the data (with random TDR parameter initialisations) and averaged *R*^2^ both over splits within each recording session (grey traces in Figure 5B) and over recording sessions (red bars in Figure 5B). We compared this to a PSTH only model in which the predicted PSTH was 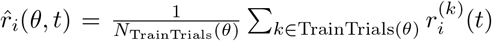.Note that a TDR model with *P* = *N*_stim_ and *w*_0,*i*_(*t*) = 0 is equivalent to this PSTH only model.

For application of TDR to classic WM models, we simulated the PSTHs (without noise) of each model under each stimulus and fit a TDR model to this set of PSTHs. We observed that a TDR model with no time component *w*_0,*i*_ was sufficient to fit these simulated PSTHs exactly, and therefore omitted this term.

Having fit the TDR model to data, we computed the Principal Component (PC) subspaces by taking the singular value decomposition of the matrix (*W*_*p*_)_*i,t*_ = *w*_*p,i*_(*t*). Writing *W*_*p*_ = *USV* ^⊺^, where *S* is a diagonal matrix containing the singular values in descending order, the columns of *U* contained the *N*-dimensional basis vectors in neural state space and the rows of *SV* ^⊺^ contained the population PSTHs. Before computing these subspaces, we first exploited several invariances of the model to transform the fitted parameters *w*_*p,i*_(*t*) and *f*_*p*_(*θ*) into a standard form (Supplementary Material, cf Aoi et al. (2020)). We plotted these population PSTHs within the top three PCs of *W*_1_ and *W*_2_, and plotted the singular value spectra (diagonal of *S*) to assess the dimensionality of each subspace *p* ∈ *{*0, 1, 2*}*.

Subspace alignments were computed using random two-way splits over trials, as for the cross-validation analysis. Given the vectors 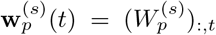 obtained from the TDR model fit to split *s*, we computed the subspace angle between the plane spanned by 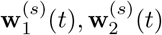 and the plane spanned by 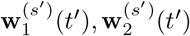 using the Matlab function *subspace*. We plotted the cosine of this subspace angle, averaged over 25 random two-way splits of the data, as a function of *t* and *t*^*′*^. The cosine of the angle between 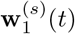 and 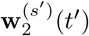 was computed in a similar fashion.

To compute the SNR of the data in the TDR Principal Component Subspace, we used a similar two-fold cross-validation split as above. First, a TDR model was fitted to the full dataset (without cross-validation). Given this fit, we formed the matrix *W*_concat_ = [*W*_1_, *W*_2_] to obtain an *N ×* 2*T* matrix describing the set of neural activity patterns generated over stimuli and time points. We performed a singular value decomposition *W*_concat_ = *USV* ^⊺^ and projected the binned spiking data onto the top six PCs as 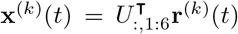. We then computed the mean and covariance of **x**^(*k*)^(*t*) over trials separately within each split of the data, conditioned on stimulus and time bin. This gave the set of linear discriminant vectors computed on split s,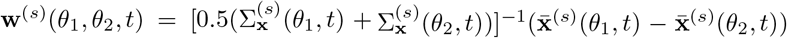. The cross-validated SNR was computed by applying the decoder trained on one split of the data to the responses on the second split, i.e. 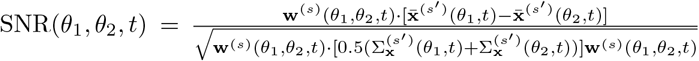. We then averaged the resulting SNR over 100 random two-way splits.

## Acknowledgements

We thank Alex Cayco-Gajic, Arthur Pellegrino, Sina Tootoonian, Matt Nolan and Matt Panichello for valuable feedback. We are particularly grateful to Matt Panichello and Tirin Moore for provision of experimental data. This work was supported in part by the Biotechnology and Biological Sciences Research Council [BB/Y513957/1].

## Supplementary Material

### A Network Model

We considered a linear dynamical system governing the evolution of the network state **x**(*t*) via the differential equation

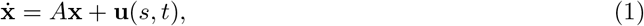

which had as parameters the connectivity matrix *A* and the stimulus-dependent input **u**(*s, t*) where the stimulus was labelled *s*. The input

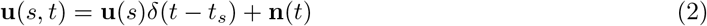

consisted of a stimulus-dependent vector **u**(*s*) indicating the direction in state space along which the network was driven by the stimulus *s* at the time of stimulus presentation *t*_*s*_, and temporally uncorrelated Gaussian white noise **n**(*t*) ∼𝒩(0, Σ_**n**_), ⟨**n**(*t*)**n**^⊺^(*t*^*′*^)⟩ = Σ_**n**_*δ*(*t − t*^*′*^). We also considered extensions to temporally extended inputs **u**(*s, t*) = **u**(*s*)ũ(*t*) + **n**(*t*) where ũ(*t*) was a boxcar function (as in Figure 1).

Given the initial condition **x**_0_ = **x**(*t*_0_), Equation 1 has the general solution for *t* ≥ *t*_0_

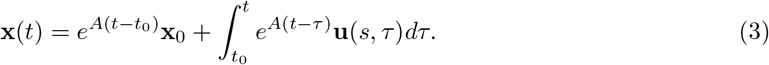

Inserting Equation 2 into Equation 3 gives (for *t, t*_*s*_ ≥ *t*_0_)

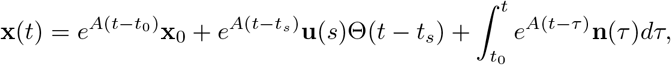

where Θ(*t*) is the Heaviside step function. Using the shorthand for the stimulus-conditioned and marginal means of a variable **a** as 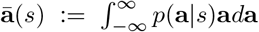 and 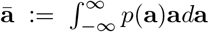, the stimulus-conditioned mean and covariance of **x**(*t*) are

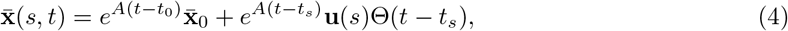

and

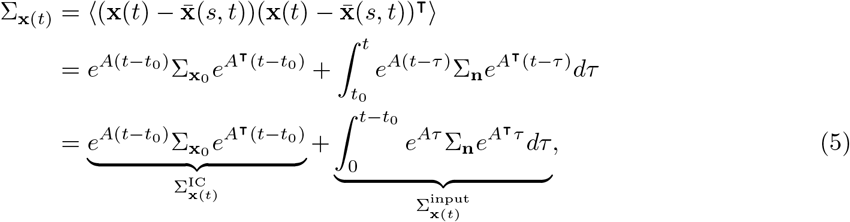

where we assumed that the initial condition is distributed as 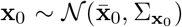 and is independent of the subsequent input noise **n** ∼ *𝒩*(**0**, Σ_**n**_). The terms 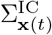 and 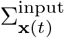 capture the response covariance generated by propagating the initial state and subsequent noise input through the recurrent dynamics, respectively. The response at time *t* under each stimulus is normally distributed over trials, with 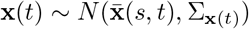, and the covariance Σ_**x**(*t*)_ is independent of the stimulus *s*.

### B Loss Functions and Their Derivatives

We consider the case where one of *M* possible input stimuli *s* ∈ *{*0, …,*M* −1*}* is presented, and optimised the performance of an optimal decoder of *s* from the network response **x**(*t*). Since the response statistics are Gaussian with stimulus-independent covariance, the performance of an optimal decoder when *M* = 2 is given by a linear readout *ŝ* = Θ(**w**_LD_ · **x**(*t*) + *c*), where 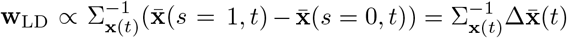 is the linear discriminant vector and *c* is a bias term (Chadwick et al., 2023). The performance of an arbitrary linear decoder **w** can be described by a signal-to-noise ratio given by:

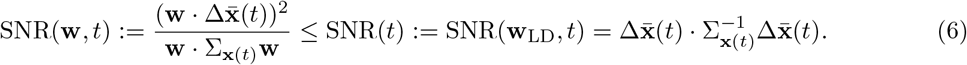

In particular, the probability of correct choice is 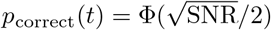, where 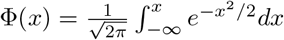 is the cumulative distribution function of the standard normal distribution (Stanislaw and Todorov,1999). Although we consider discrete classification tasks, the SNR also places a lower bound on the decoding error for continuous estimation tasks. In particular, the Fisher information is given by 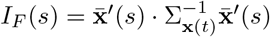, and the Cramer-Rao lower bound is ⟨(*s* − *ŝ*)^2^⟩ ≥ 1*/I*_*F*_ (*s*).

For two-stimulus tasks, we therefore defined the loss function 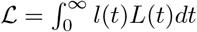, where 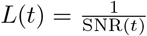 and *l*(*t*) is an arbitrary weighting function. We chose to minimise the reciprocal of the SNR rather than directly maximising the SNR, as this avoids solutions where a low SNR at time point *t* is compensated by a high SNR at *t*^*′*^.

The derivative of this loss function with respect to the weights is

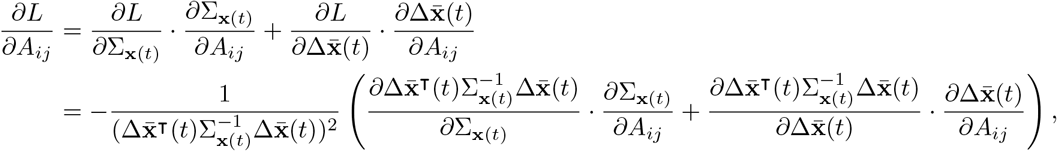

where *M* · *N* = Trace(*MN*^⊺^) = Σ_*i,j*_ *M*_*ij*_*N*_*ij*_ denotes a sum over elements. Given the derivatives 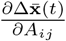 and 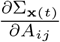, the above equation allows for a direct computation of the gradient of the loss function with respect to *A*. We describe how these derivatives can be computed in Section C.

For the general setting of networks with *N* neurons, *M* stimuli, and a penalty on response energy, we optimised the loss

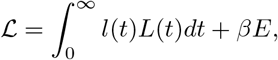

where 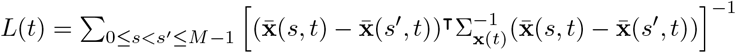 and 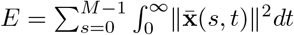.

### C. Derivatives of the Network Statistics with Respect to the Weights

To perform gradient descent on this loss function, we computed the derivatives of the mean 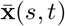, covariance Σ_**x**(*t*)_ and energy *E* with respect to the weight matrix *A*.

#### Derivative of mean response

The derivative of the mean response (Equation 4) is

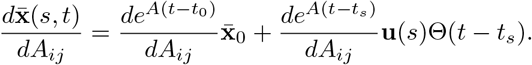

The loss functions we consider depend on the difference in mean response to the two stimuli 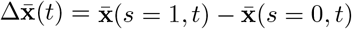, which has derivative:

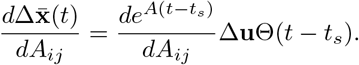

The derivative of the matrix exponential can be computed numerically (Najfield and Havel, 1995; Al-Mohy and Higham, 2008), and a solver is available via the `expm_frechet` function in SciPy’s Linear Algebra library.

#### Derivative of response covariance

The derivative of the covariance (Equation 5) requires the solution of two Lyapunov equations, with numerical solvers widely available. The first Lyapunov equation is the result of integrating 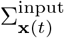 by parts

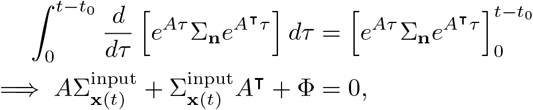

where 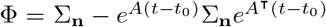. The second Lyapunov equation results from differentiating the first with respect to *A*_*ij*_ to obtain

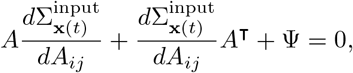

where 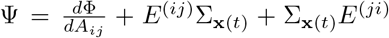, with 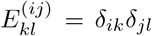 and 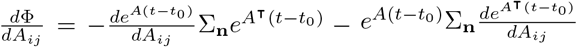. Since this is another Lyapunov equation it can be solved in terms of *A* and Ψ to obtain 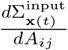. The derivative of the network covariance 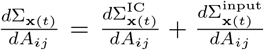 can then be evaluated by handling 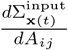 as described here and 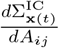 using a numerical solver for the derivative of the matrix exponential, as described above for the derivative of the mean (assuming that the initial state covariance 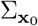 is independent of the weights *A*).

#### Limiting cases for response covariance

Two special cases allowed us to neglect the role of the initial condition-driven covariance 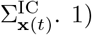 the scenario where the network was initialised at stationary state with respect to the input noise Σ_**n**_, which can be found by setting *t*_0_ → −∞, so that Φ = Σ_**n**_ and 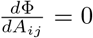, and 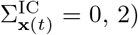 the scenario where 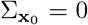, which corresponds to a fixed (zero-variance) initial condition. We focused on these special cases in order to reduce the space of free parameters in the model and to gain insight into the behaviour of the model in these limiting situations.

#### Derivative of the network energy term

The energy *E* and its gradient 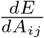 could be computed directly without numerical integration using 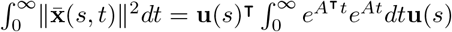, with the inte-gral and its derivative having direct solutions via Lyapunov equations, as was the case for the network covariance.

### D Relationship to Backpropagation and Other Learning Rules

The loss function and optimisation procedure described above are based on the trial-averaged statistics of the network response. How does this relate to learning rules that operate based on trial-to-trial feedback, such as reward-based Hebbian learning, backpropagation through time, or reinforcement learning? To address this question, we show that 1) in the limit of slow learning rate, any learning rule that maximises the fraction of correct choices based on trial-by-trial feedback will converge to the same weight updates as our method 2) in the limit of large batch size, backpropagation through time on a squared error loss function produces similar weight updates to our method and converges to the same solution.

#### Relationship to learning from trial-by-trial feedback

The objective we optimise *L* = 1*/*SNR is directly related to the probability of a correct decision *p*_correct_ using an optimal decoder, since 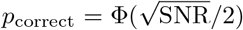 where 0.5 ≤ Φ(*x*) ≤ 1 is monotonically increasing (as described above). Thus, 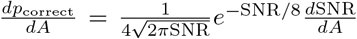 and 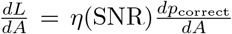. The factor 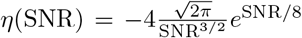 depends only on the SNR, so that the two gradients differ only up to a scaling factor, and therefore produce identical weight trajectories. In the limit of a small learning rate, any learning rule that optimises the fraction of correct choices using trial-by-trial feedback will converge to the gradient 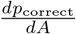, and will therefore follow the same learning dynamics as our method.

#### Relationship to backpropagation through time

Methods which do not explicitly optimise the probability of correct choice may nevertheless yield very similar learning dynamics to those produced by our method. To illustrate this finding, we derive analytical expressions for the updates generated via backpropagation through time (BPTT) and show that numerical implementation of these updates generates near-identical weight trajectories to those produced by our method. Specifically, we consider BPTT on the continuous-time linear recurrent neural network 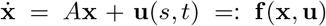 with **u**(*s, t*) = **u**(*s*)*δ*(*t* − *t*_*s*_) + **n**(*t*). On each trial, we consider a trajectory from initial time point *t*_0_ to decision time *t*_*d*_ with a loss *L* that depends only on the final time point **x**(*t*_*d*_). The continuous-time BPTT updates are then given by

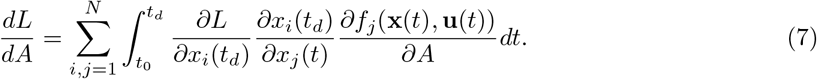

We consider the squared error loss function *L*(*t*_*d*_) = (**w** · **x**(*t*_*d*_) + *c* − *s*)^2^ = (*ŝ* − *s*)^2^. The terms in Equation (7) are the 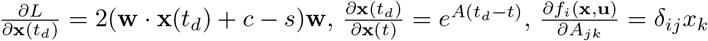. Therefore

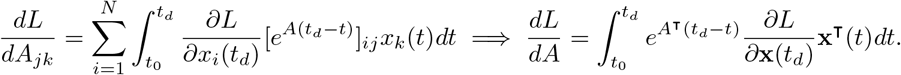

We next consider the stationary state limit *t*_0_ → −∞, with

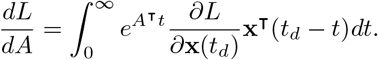

Inserting the expressions derived above for 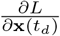 and 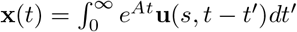 we obtain

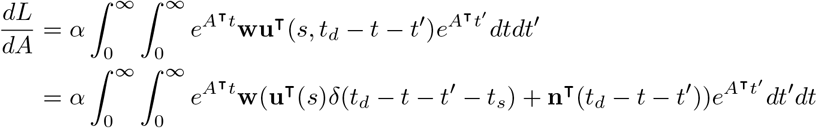

where 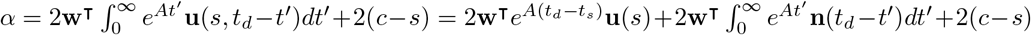.Using ⟨**n**(*t*)⟩ = 0 and ⟨**n**(*t*)**n**^⊺^(*t*^*′*^)⟩ = Σ_**n**_ *δ*(*t*− *t*^*′*^), we then compute the expected gradient over all possible realisations of the noise term **n** (for fixed *s*):

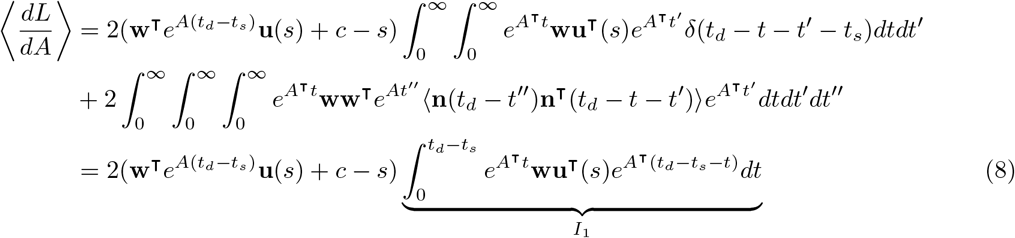

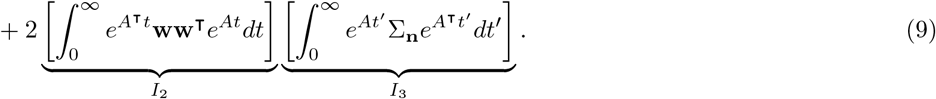

Each of the above integrals *I*_1_, *I*_2_, *I*_3_ can be reformulated as a Lyapunov or Sylvester equation, with numerically efficient and stable solvers available. In particular, *I*_2_ = lyap(*A*^⊺^, **ww**^⊺^) and *I*_3_ = lyap(*A*, Σ_**n**_). Integrating *I*_1_ by parts gives 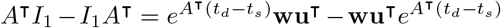, which is a Sylvester equation of the form *PI*_1_ + *I*_1_*Q* = *C*. However, this Sylvester equation does not have a unique solution^1^. Instead, a unique solution for *I*_1_ can be found by taking the eigendecomposition 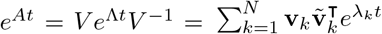, which gives

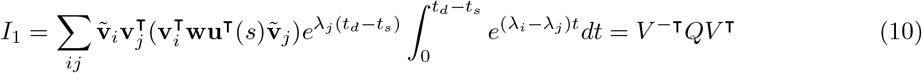

where *Q* = (*V* ^⊺^**wu**^⊺^*V* ^*−*⊺^) ⊙ *S*, with 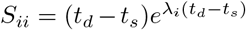 and 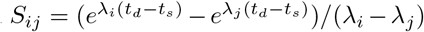 (for *i* ≠ *j*), with ⊙ the Hadamard (or element-wise) product. Thus, each update of *A* during the BPTT algorithm can be achieved by solving two Lyapunov equations (for *I*_2_ and *I*_3_) and performing an eigendecomposition on *A* to compute *I*_1_.

To complete the BPTT algorithm, we require updates for the readout weights **w** and threshold *c*. These updates are given as follows:

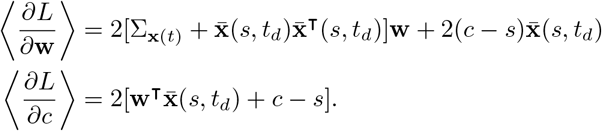

These readout parameters could be updated via gradient descent alongside the recurrent weights *A*. However, if we assume that the updates of the recurrent weights are slow relative to the readout weights, we can set

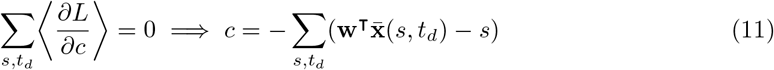

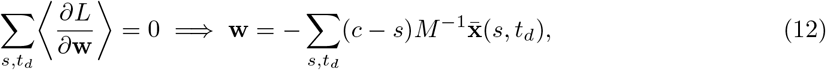

where 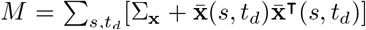, and we included a sum over time points to handle the case of a fixed (time-independent) readout over a range of delays. Combining these two expressions gives:

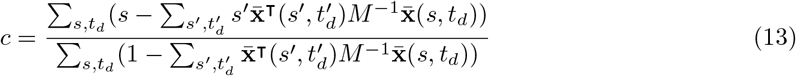

with **w** obtained by inserting this expression into Equation (12). Taken together, the full BPTT algorithm involves iteratively using Equations (12-13) to update readout weights followed by Equations (8-10) to update the recurrent weights (summed over stimuli and time points). An implementation of this algorithm is shown in Supplementary Figure S4.

### E Optimal Solution to Task: An Ideal Observer Analysis

We consider an ideal observer of the network input **u**(*s, t*) = **u**(*s*)ũ(*t*) + **n**(*t*) tasked with reporting the stimulus identity *s* ∈ {0, 1} at time *t*_*d*_. Assuming that the input noise is Gaussian, stimulus-independent, and temporally uncorrelated, the optimal solution involves two steps: 1) projecting the inputs onto the linear discriminant vector 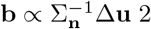) applying a temporal filter *f* that selects and delays inputs over time. Here, we formalise this solution mathematically, showing that the optimal solution is a form of matched filtering (Turin, 1960).

We consider a linear projection and temporal filtering of the input time series **u**(*s, t* ≤ *t*_*d*_) onto a vector **w** and filter 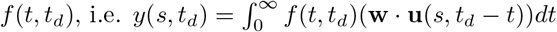 with

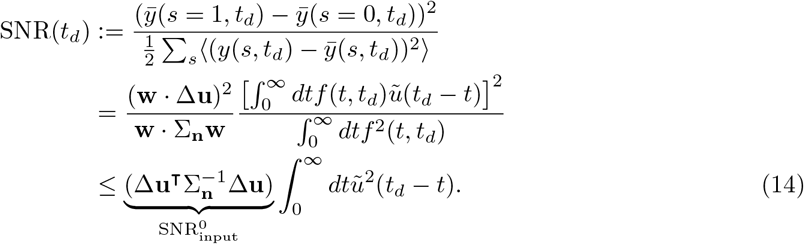

In particular, the above SNR is maximised when 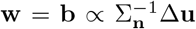 and *f*(*t, t*_*d*_) ∝ ũ(*t*_*d*_ *− t*). These filtering and projection operations can be understood as linear discriminant analysis over both neurons and time, and can also be derived as the maximum likelihood estimate of *s* given the full input time series up to time *t*_*d*_, i.e. by maximising *p*(**u**(*s, t* ≤ *t*_*d*_) *s*), with 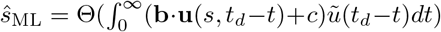. Thus, Equation (14) quantifies the performance of a Bayes optimal ideal observer of the network input (assuming equal prior likelihood of each stimulus).

The above solution requires a filter that depends on the optimised readout time, also known as a time-varying linear filter. In contrast, a fixed filter optimised for a single readout time *t*_*d*_ will have a suboptimal output SNR for *t* ≠ *t*_*d*_

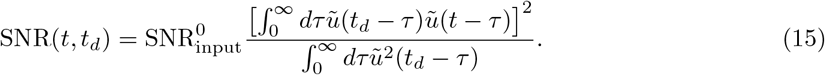

To illustrate these solutions, we provide examples from three tasks: first, a working memory task with instantaneous inputs, ũ_*δ*_(*t*) = *δ*(*t* − *t*_*s*_); second, an evidence integration task where ũ_Θ_(*t*) = Θ(*t* − *t*_*s*_); third, a working memory task where the stimulus appears for a fixed interval of time, modelled as a “boxcar” function ũ_∏_(*t*) = Θ(*t* − *t*_*s*_) − Θ(*t* − (*t*_*s*_ + *T*)). The optimal time-varying filters for these tasks are *f*_*δ*_(*t, t*_*d*_) = *δ*(*t*_*d*_ − *t*_*s*_ − *t*), *f*_Θ_(*t, t*_*d*_) = Θ(*t*_*d*_ − *t*_*s*_ − *t*), *f*_∏_(*t, t*_*d*_) = Θ(*t*_*d*_ − *t*_*s*_ − *t*) − Θ(*t*_*d*_ − *t*_*s*_ − *T* − *t*).

Following Equation (14), these filters give rise to the following SNRs:

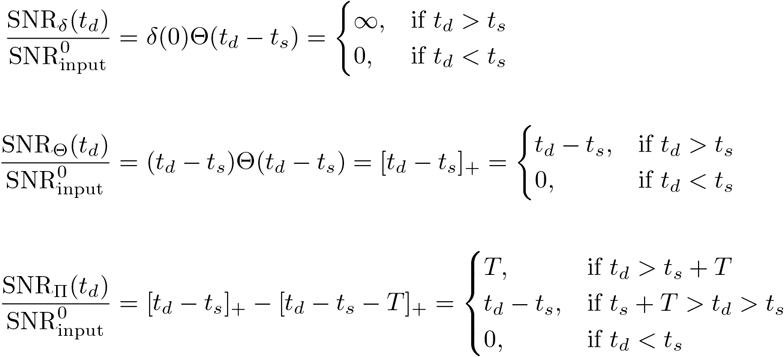

These solutions are illustrated in Supplementary Figure S2, where we also show the performance of fixed filters optimised for a single readout time (Equation (15)).

### F Analysis of Linear Dynamical System Task Performance

We next sought to determine the performance of linear network models on the working memory task. The above ideal observer analysis provides an upper bound on the SNR of network responses, and suggests that networks could be suboptimal due to two factors: 1) a failure to correctly combine inputs to different neurons via a suboptimal projection **w**, or 2) a failure to correctly combine inputs across time via the wrong temporal filtering *f*.

We show that attractor networks (those with orthogonal eigenvectors, also called normal networks) perform poorly on working memory tasks, with their SNR decaying at best as 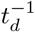. We then show that (non-normal) functionally-feedforward networks can approximate the ideal observer solution to an arbitrary level of precision, but require exponential increases in firing rates to achieve this. Finally, we show that networks with rotational dynamics achieve higher SNR than both attractor and feedforward networks.

In the following we consider networks receiving delta pulse inputs 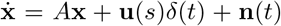 (we set *t*_*s*_ = 0 for convenience). We consider the signal-to-noise ratio of network responses along a readout vector **w** defined as 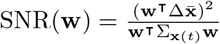. We will often consider the optimal readout 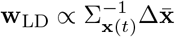 with 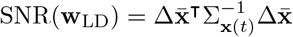.

#### F.1 SNR Along Individual Dynamical Modes

Previous work has suggested that alignment of dynamical modes with the input linear discriminant provides a neural mechanism for working memory and evidence integration tasks (Seung, 1996; Mante et al., 2013; Chadwick et al., 2023; Stroud et al., 2023; Pagan et al., 2025). We therefore first investigated the SNR along individual dynamical modes of a network.

To decompose the dynamics into individual modes, we use the eigendecomposition 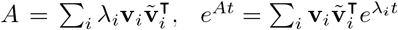, where 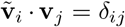 are left and right eigenvectors chosen such that ∥**v**∥ ^*i*^ = 1. Given Equations (4) and (5), the mean and variance of network output projected onto a left eigenvector are (for *t* ≥ *t*_*s*_ = 0 and *t*_0_ ≤ *t*_*s*_ = 0)

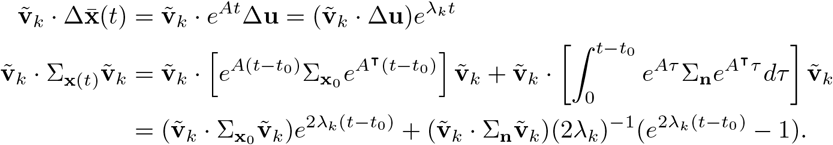

Thus,

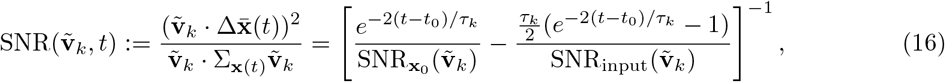

where 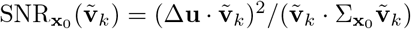 and 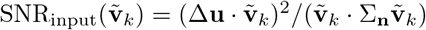.

#### Optimal dynamics at stationary state

If the network is initialised at stationary state with respect to the input noise (i.e. if *t*_0_ → −∞), the response SNR is

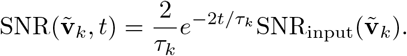

This solution is optimised when the left eigenvector is aligned to the input discriminant, i.e. 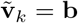. The SNR depends non-monotonically on the integration time constant *τ*_*k*_ = −1*/λ*_*k*_, and has an optimal time constant for readout time *t*_*d*_ given by *τ*_*k*_ = 2*t*_*d*_. Thus, a single eigenmode is optimised for a readout at time *t*_*d*_ if its left eigenvector is aligned to the input discriminant and its eigenvalue is 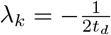. However, this optimal SNR decreases as a function of the optimised readout time according to SNR(**b**, *t*_*d*_) = (*et*_*d*_)^*−*1^SNR_input_(**b**). This is in contrast to the optimal ideal observer solution, whose performance is independent of the readout time provided *t*_*d*_ ≥ *t*_*s*_.

#### Optimal dynamics for fixed initial state

If the network is initialised at a fixed (i.e. zero-variance) initial condition **x**_0_ at time *t*_0_ ≤ *t*_*s*_ = 0, the response SNR is:

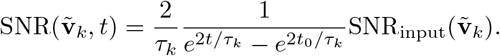

Plotting this equation numerically as a function of *λ*_*k*_ = −1*/τ*_*k*_ for different values of *t*_0_ shows that stable dynamics (*λ*_*k*_ *<* 0) are optimal when |*t*_0_| *> t*, neutrally stable dynamics (*λ*_*k*_ = 0) are optimal when |*t*_0_| = *t*, and unstable dynamics (*λ*_*k*_ *>* 0) are optimal when |*t*_0_| *< t* (Supplementary Figure S6B-D). The optimal *λ*_*k*_ scales monotonically with *t*_0_, respectively approaching infinitely strong amplification *λ*_*k*_ → ∞ as *t*_0_ → 0 and the stationary state solution 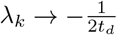 as *t*_0_ → −∞. The amplifying dynamics for |*t*_0_| *< t* act to amplify the signal throughout the delay, while the decaying dynamics for |*t*_0_| *> t* are required to avoid amplification of pre-stimulus noise input above the signal.

These solutions are highly suboptimal with respect to the ideal observer solution. This suboptimality arises because an individual eigenmode implements an exponential filter with time constant *τ*_*k*_ = *−*1*/λ*_*k*_, and therefore integrates noise before and after stimulus presentation. The ideal observer solution has a narrow integration time window but a long memory time constant, and departs substantially from a simple exponential filter.

#### F.2 Attractor Networks

For attractor (normal) networks, the eigenvectors form an orthogonal basis with 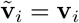 and **v**_*i*_ · **v**_*j*_ = *δ*_*ij*_. As a consequence, their response SNR follows trivially from the above analysis of individual eigenmodes as we show below. The mean response of an attractor network to the inputs considered above is

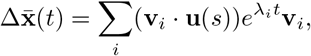

and the response covariance is

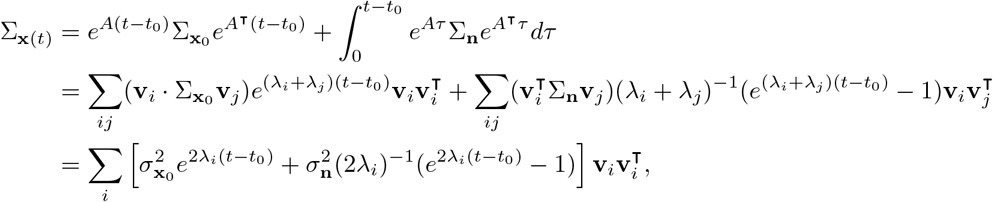

where the last step assumed that 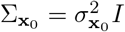 and 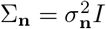.

The SNR of network output at time *t* is

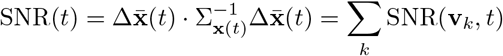

where SNR(**v**_*k*_, *t*) is the single-mode SNR given in equation (16). The SNR of an attractor network is therefore maximised when a single eigenmode is aligned to the input discriminant, i.e. **v**_*k*_ = **b**, in which case the solution reduces to that of single modes considered above, with all other modes having zero SNR.

Thus, attractor networks perform poorly on the working memory task, with SNR decaying exponentially over time due to noise (and decaying as 1*/t*_*d*_ as a function of the optimised delay). We next show how functionally-feedforward (non-normal) and rotational dynamics can improve upon attractor mechanisms.

#### F.3 Preliminary: The Real and Complex Schur Decomposition

While the eigendecomposition of the matrix *A* is sufficient to characterise the dynamics and task-performance of a normal network, this description often obscures the transient dynamics of non-normal and rotational networks (Ganguli et al., 2008; Goldman, 2009; Murphy and Miller, 2009). Instead we turned to the Schur decomposition to characterise such networks. Here we briefly describe the real and complex forms of the Schur decomposition that we use extensively in the following sections.

##### Complex Schur Decomposition

Any matrix *A* ∈ *ℝ*^*N ×N*^ can be expressed in Schur form as *A* = *ULU* ^*†*^ where *U* ∈ *ℂ*^*N ×N*^ is a unitary matrix, i.e. *U* ^*−*1^ = *U* ^*†*^ where *U* ^*†*^ := (*U*^⊺^)^*∗*^, and *L* ∈ *ℂ*^*N ×N*^ is an upper-triangular matrix, i.e. *L*_*i,i*+*k*_ = 0 for all *k <* 0. Given a connectivity matrix *A*, the off-diagonal elements of *L* describe “functionally-feedforward” weights between a set of orthogonal neural activity patterns (Schur modes, i.e. columns of *U*). However, when *A* has complex eigenvalues, the elements of both *U* and *L* become complex-valued, which complicates this interpretation.

##### Real Schur Decomposition

When some eigenvalues are complex, a more interpretable basis in which to analyse the dynamics is given by the real Schur decomposition *A* = *QT Q*^⊺^, where *Q* and *T* are real-valued matrices, with *Q*^*−*1^ = *Q*^⊺^ (i.e., *Q* is an orthogonal matrix) and *T* a block upper-triangular matrix

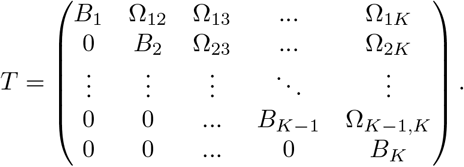

Here, *K* is the number of real eigenvalues plus the number of pairs of complex conjugate eigenvalues and the diagonal blocks *B*_*i*_ are either 1*×*1 submatrices corresponding to real eigenvalues of *A* (i.e., *B*_*i*_ = *λ*_*k*_) or 2 *×* 2 submatrices corresponding to complex conjugate eigenvalue pairs (with 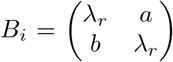 for some *a, b* ∈ *ℝ* such that the eigenvalues of *B*_*i*_ are equal to a pair of complex-conjugate eigenvalues of *A, λ* = *λ*_*r*_ *± iλ*_*ω*_). Since the columns of *Q* are orthonormal, the real Schur decomposition provides an orthonormal basis partitioned into one-dimensional non-rotational modes and two-dimensional rotational planes, with functionally-feedforward interactions between these one- and two-dimensional subspaces given by the off-diagonal blocks Ω_*ij*_. Thus, the Ω_*ij*_’s are either 1 *×* 1, 2 *×* 1, 1 *×* 2, or 2 *×* 2 matrices capturing functionally-feedforward interactions between these modes.

##### Two-Dimensional Case

A matrix with eigendecomposition 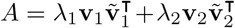 and *λ* ∈ *ℝ* can be expressed in Schur form as 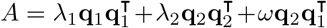, where *ω* is a functionally-feedforward weight from the input Schur mode 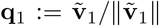 to the output Schur mode **q**_2_ := **v**_2_. Thus, *Q* = [**q**_2_, **q**_1_] and

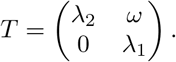

A matrix *A* ∈ ℝ^2^ with complex eigenvalues *λ*_*r*_ *± iλ*_*ω*_ can be written in real Schur form as *A* = *QT Q*^⊺^ where

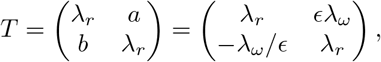

with 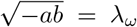 and *Q*^⊺^ = *Q*^*−*1^ ∈ *ℝ*^2*×*2^ a real orthogonal matrix. In the second equality, we introduced the parameter *ϵ* = *a/λ*_*ω*_, which can be understood as the ellipticity (or eccentricity) of the rotation, and is closely related to the non-normality of the system. In this basis, the major and minor axes of the ellipse are aligned to the two basis axes (defined by the columns of *Q*). Thus, in real Schur form the matrix *A* is characterised by four parameters: the real and imaginary parts of the eigenvalues, *λ*_*r*_ and *λ*_*ω*_, which characterise the damping rate and rotational frequency respectively; the ellipticity of the orbit *ϵ*; and the angle of the major axis of the ellipse in neural state space (determined by the orthogonal matrix *Q*). The matrix exponential of *A* is *e*^*At*^ = *Qe*^*Tt*^ *Q*^⊺^, and the matrix exponential of *T* can be computed directly as

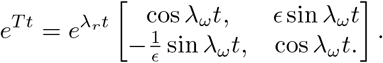

Importantly, these results for two-dimensional systems also apply to any 2 × 2 submatrix *B*_*i*_ in the *N*-dimensional real Schur decomposition.

##### Invariances of the Real Schur Decomposition

The Schur decomposition is non-unique: each ordering of eigenvalues leads to a different *Q* and *T*. For the real Schur decomposition, this can be factored into an ordering of blocks and an ordering within 2 × 2 blocks. Within a 2 × 2 block, the ordering of eigenvalues makes only a superficial difference to the resulting decomposition. In particular, if *k, k* + 1 corresponds to a rotational plane (i.e., if *T*_*k*:*k*+1,*k*:*k*+1_ = *B*_*i*_ for some *i*), the change of Schur basis *Q* = [**q**_1_, **q**_2_, …, **q**_*k*_, **q**_*k*+1_, …, **q**_*N*_] → [**q**_1_, **q**_2_, …, **q**_*k*+1_, **q**_*k*_, …, **q**_*N*_] and *T*_:,*k*_ ↔ *T*_:,*k*+1_, *T*_*k*,:_ ↔ *T*_*k*+1,:_ yields another real Schur decomposition with *ϵ* → −1*/ϵ*. A further invariance for both real and complex Schur decompositions is to reverse the sign of a mode, i.e. *Q* = [**q**_1_, **q**_2_, …, **q**_*k*_, …, **q**_*N*_] → [**q**_1_, **q**_2_, …, **q**_*k*_, …, **q**_*N*_], *T*_*k*,:_ → −*T*_*k*,:_, *T*_:,*k*_ → −*T*_:,*k*_ with *ϵ* → −*ϵ*. Through this combination of reordering and sign-reversal of real Schur modes, it is possible to select a real Schur decomposition for which rotations are clockwise or anticlockwise (*ϵ* ≷ 0) and have their major axis aligned to the x- or y-axis (|*ϵ*| ≷ 1), and have either positive or negative loading onto the input discriminant (**q**_*k*_ · **b** ≷ 0). For numerically optimised networks, we utilised these invariances to define a unique Schur decomposition in which 1) the decay time constants *τ* = −1*/λ*_*r*_ are in descending order from input to output mode 2) elliptical orbits have their major axis aligned to the x axis (|*ϵ*| *>* 1) 3) rotations in each plane are clockwise (*ϵ >* 0) 4) the input mode to each plane has positive alignment with the input discriminant.

#### F.4 Two-Dimensional Feedforward Networks

We first considered the performance of feedforward (non-normal) networks with real eigenvalues on the working memory task. We restrict our analysis to the stationary state limit *t*_0_ → −∞. Using the Schur basis discussed in Section F.3, the dynamics of a two-dimensional non-normal network with real eigenvalues decouples into two equations:

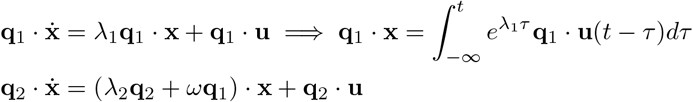

The mean and variance of responses along the first mode are given by the single-mode analysis in Section F.1:

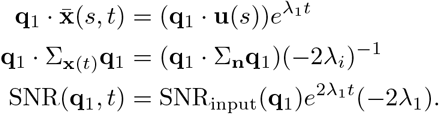

For the second mode these are given by

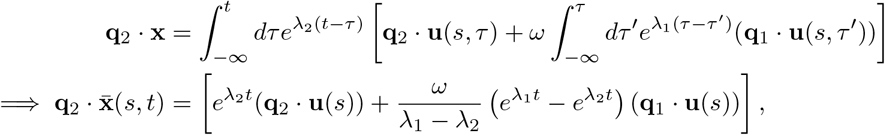

and

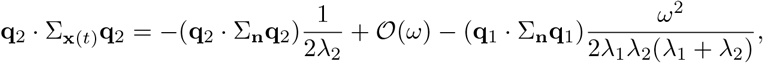

where we omitted the explicit form of the 𝒪(*ω*) interaction term for brevity. In the non-normal limit *ω* → ∞, the response SNR along the output Schur mode becomes:

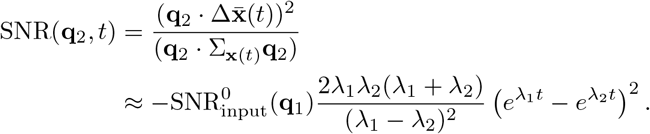

This SNR behaves non-monotonically as a function of time, reaching a peak at a finite time before decaying back to zero. This non-monotonic timecourse is due to the transfer of information from the input to output mode over time, and the fact that the readout is restricted to the output mode - the SNR of the optimal (time-varying) readout decreases monotonically due to a shift in weighting of the readout from input to output mode over time (Section F.6).

For a given choice of eigenvalues, the SNR is maximal at the delay *t*_*d*_ = log(*λ*_2_*/λ*_1_)*/*(*λ*_1_ − *λ*_2_). How should the eigenvalues then be chosen in order to optimise response SNR at a particular delay *t*_*d*_? Writing *λ*_1_ = *λ, λ*_2_ = *λ* − *δλ* and taking the limit *δλ* → 0, we have *t*_*d*_ = log(1 − *δλ/λ*)*/δλ* ≈ −1*/λ* and

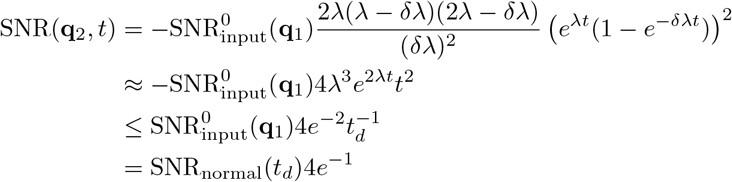

where the inequality gives the behaviour of a network with *λ* optimised for the delay *t*_*d*_, and 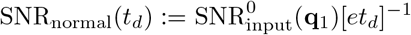 is the SNR of a normal network optimised for readout time *t*_*d*_. Compared to the optimal solution of the normal network with stationary state covariance, the response SNR in this non-normal solution is scaled by a factor of 4*e*^*−*1^ ≈ 1.47. We note the limits *ω* → ∞ and *λ*_1_ ≈ *λ*_2_ taken above are consistent with the observed behaviour of networks during optimisation before transition to rotational dynamics (Supplementary Figure S3E-F).

#### F.5 Two-Dimensional Rotational Networks

We next considered a two-dimensional rotational network for which the two eigenvalues and eigenvectors of the weight matrix *A* form a complex conjugate pair *λ, λ*^*∗*^ and **v, v**^*∗*^ (where **x**^*∗*^ = Re(**x**) −*i*Im(**x**) is complex conjugation). Using the results of Section F.3, the mean and stationary state covariance of the network response can be computed in the real Schur basis as

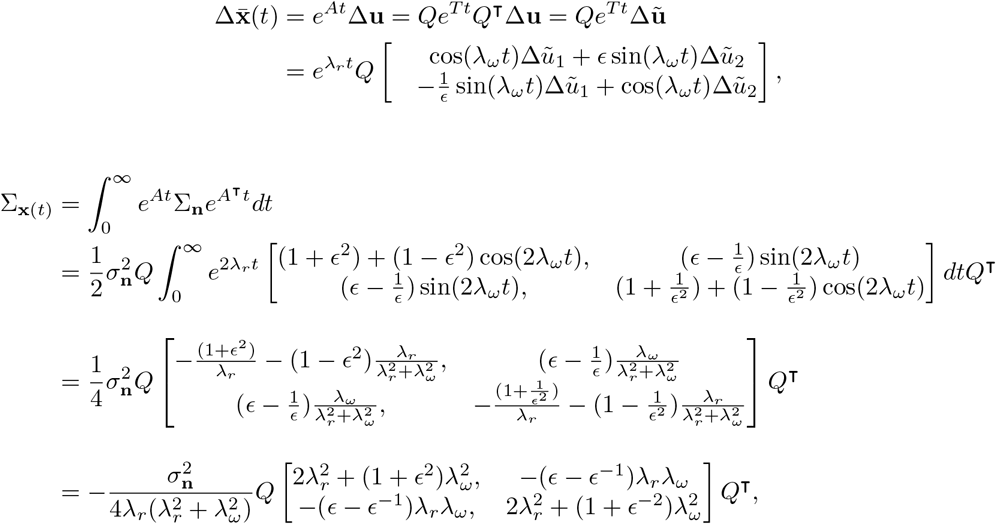

where we defined **ũ** = *Q*^⊺^**u** and assumed that 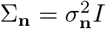.

The covariance matrix can be inverted directly as

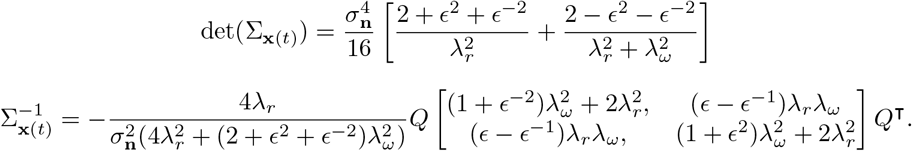

The SNR of the rotational network takes on a simple form when the input linear discriminant is aligned to the one of the Schur modes. In particular, setting Δũ_1_ = 0, the SNR is

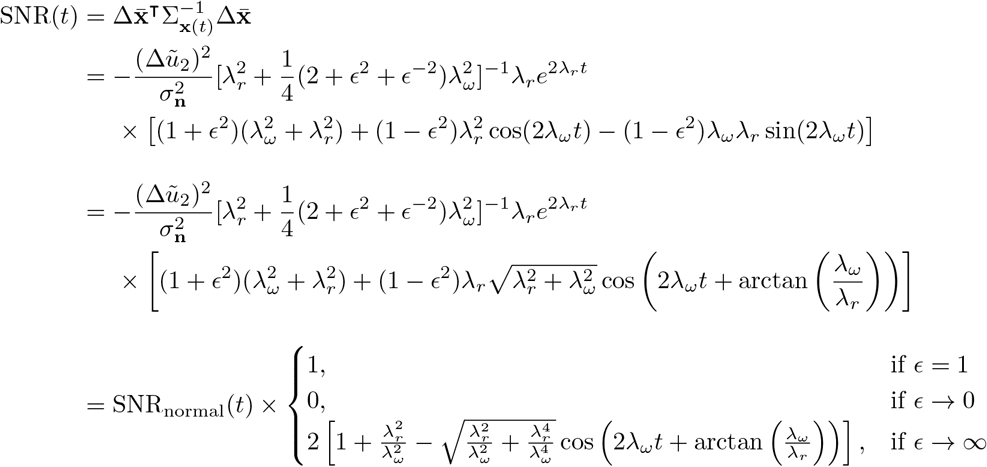

The SNR when *ϵ* = 1 is identical to that of a normal network with real eigenvalue *λ* = *λ*_*r*_ (given by 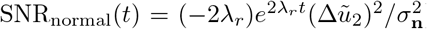), independent of the rotational frequency *λ*_*ω*_. When *ϵ* → *∞*, the SNR derived above matches that observed in optimised networks (Supplementary Figure S8B). This case corresponds to a highly elliptical rotation, with the input linear discriminant aligned to the minor axis of the ellipse. Indeed, when optimising numerically (without an energy penalty), we found that *ϵ* → *∞* with increasing numbers of iterations, albeit very slowly and with only marginally increasing SNR. The third limit, *ϵ* → 0, corresponds to a highly elliptical rotation with the input discriminant aligned to the major axis of the ellipse. In this case, noise input to the minor axis of the ellipse is strongly amplified, but the input signal is damped, leading to low SNR.

#### F.6 N-Dimensional Functionally-Feedforward Networks

We next considered delay line architectures, in which multiple feedforward modes are chained together in sequence (Ganguli et al., 2008; Goldman, 2009). Previous studies have shown that such architectures maximise the SNR of discrete-time linear dynamical systems, provided the number of neurons is equal to or greater than the number of time steps in the delay (Ganguli et al., 2008). However, the optimal dynamics for continuous-time networks, or for discrete-time networks with more time steps than neurons, was not determined. While Goldman (2009) considered a continuous-time network and showed that a delay line architecture can generate responses with memory time constant proportional to the number of neurons, they did not consider noise robustness or show that such architectures are optimal. We derive analytical expressions for the signal-to-noise ratio of a continuous-time delay line and show that, although SNR scales with the number of neurons, it is outperformed by the rotational solutions obtained via numerical optimisation (Supplementary Figure S1, Figure 3F).

##### F.6.1 Mean and Covariance of Network Response

We considered networks with *A*_*ij*_ = *λδ*_*ij*_ + *ωδ*_*i,j−*1_^2^. This network is known as a homogeneous delay line Ganguli et al. (2008). In this basis, the network is structurally feedforward with each element *i* corresponding to a single neuron. However, a rotation of the connectivity matrix *A* → *QAQ*^⊺^ produces a functionally-feedforward network for which the feedforward structure acts on Schur modes rather than single neurons.

The matrix exponential for the homogeneous delay line can be computed directly from its power series, 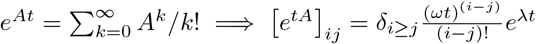, where *δ*_*i≥j*_ is 1 for *i* ≥ *j* and zero otherwise. If the network is initialised in the state *x*_*i*_(0) = *δ*_*i*1_, then in the absence of input the response propagates down the delay line as 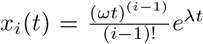. Thus, if the stimulus inputs **u**(*s, t*) = **u**(*s*)*δ*(*t*) + **n**(*t*) are such that Δ*u* = ∥Δ**u**∥*δ*_*i*1_, the trial-averaged responses follow 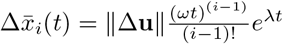. The station-ary state covariance matrix for the delay line driven by Gaussian white noise input 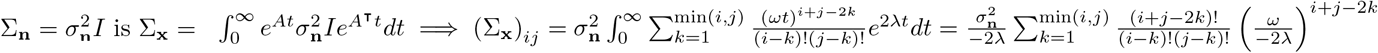.

##### F.6.2 Response SNR of Individual Schur Modes

In this section, we place a tractable lower bound on the total response SNR by computing the SNR of the *i*th Schur mode. Writing 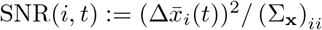, we have

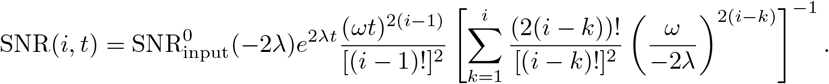

In the limit *ω* → ∞ the first term in the sum dominates, giving

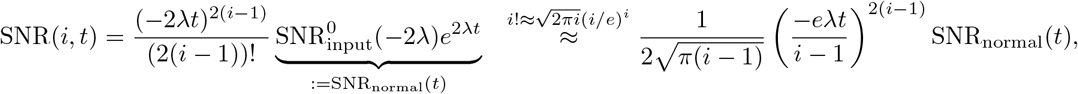

where SNR_normal_(*t*) is the SNR of a normal network with a left eigenvector aligned to the input discriminant and Stirling’s approximation was used for *i* ≫ 1. This SNR is maximised for a delay *t*_*d*_ when *λ* = −(*i* − 1*/*2)*/t*_*d*_, with

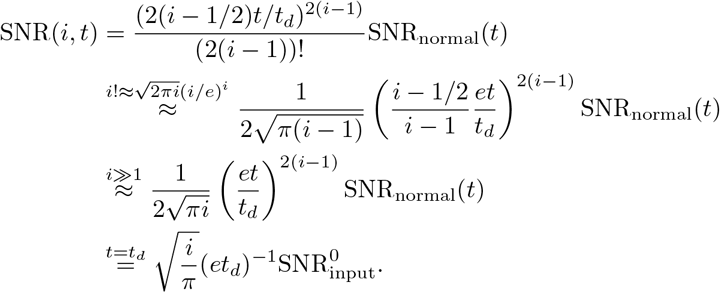

Thus, although the optimal response SNR decays as 1*/t*_*d*_, it also grows with the number of neurons as 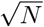, and an arbitrarily high SNR can be achieved for any delay *t*_*d*_ by increasing the length of the delay line. Note that the optimal time constant decreases as a function of the length of the delay line as as *τ* = −1*/λ* ∼ *t*_*d*_*/N*.

##### F.6.4 Response SNR of Optimal Population Readout

The previous section considered the performance of a single-mode readout, which is in general suboptimal. We next derive a general expression for the SNR of the optimal readout of the delay line, and calculate this SNR explicitly for the two-dimensional case.

The SNR of the network response at time *t* along a readout **w** is

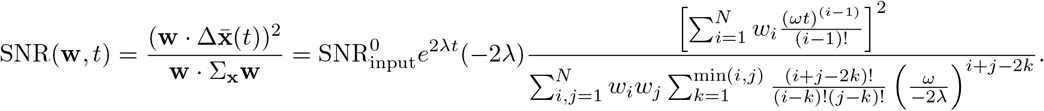

This can be written as:

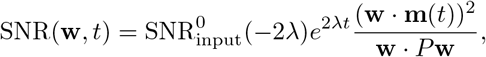

where 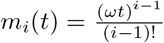 and 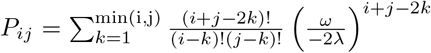. The SNR is maximised using the linear discriminant readout **w** ∝ *P* ^*−*1^**m**(*t*), which gives

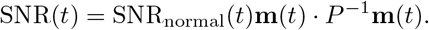

Thus, the factor **m**(*t*) · *P* ^*−*1^**m**(*t*) measures the gain in performance introduced by non-normal integration.

To determine the behaviour of this SNR in the large *ω* limit, we define 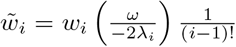, which gives

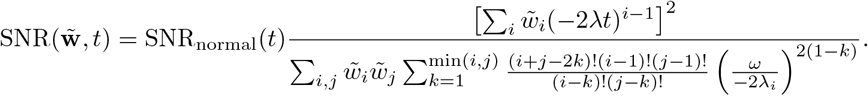

In the limit *ω* → ∞, this becomes

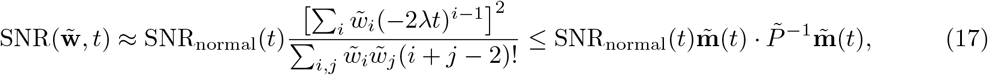

where 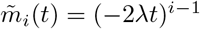 and 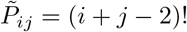 and equality is achieved by setting 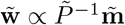.

##### Two-Dimensional Case

For a two-dimensional network, 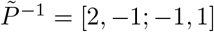, so that 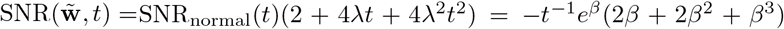 where *β* = 2*λt* and we set 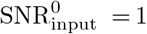. The extrema of this SNR for a given delay *t*_*d*_ are found by solving 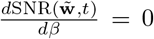, which gives *β*^3^ + 4*β*^2^ + 4*β* + 2 = 0. This equation has three real roots at 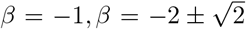. The first root (*β* = 1) has 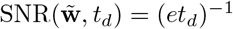, which is equal to the SNR of a normal network optimised for *t*_*d*_. The second and third roots 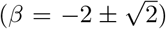 have 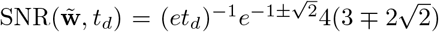. The root 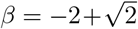 has 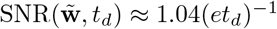, while the root 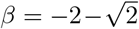 has 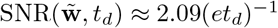.

We numerically plot the solution to Equation (17) for different values of *N* (with optimal *λ*) in Figure 3F), finding an approximately linear scaling of response SNR with the number of neurons *N*.

#### F.7 Encoding Multidimensional Stimulus Spaces Using Orthogonal Subspaces

For tasks in which multiple stimuli *s* ∈ *{s*_0_, *s*_2_, …, *s*_*M−*1_*}* can be presented, we considered the SNR for each stimulus pair SNR(*s*_*i*_, *s*_*j*_, *t*). Here, we show how networks optimised for discrimination of two stimuli can naturally be composed to form networks that discriminate arbitrarily many stimuli within a *P*-dimensional space with SNR for any stimulus pair equal to that of an *N*-dimensional network optimised exclusively for that stimulus pair. The assumption that inputs span a linear subspace is true in particular for the case of cosine-tuned inputs considered in the main text (where *P* = 2), but we present results for the general case here.

− ∝ ∈

To construct such a network, we start with an *N*-dimensional network optimised for discrimination of two stimuli *s*_1_, *s*_2_, with *A* = *QT Q*^⊺^ and **u**(*s*_2_) − **u**(*s*_1_) **b ℝ** ^*N*^. We then use this network to construct a network that discriminates a set of input stimuli that span a *P*-dimensional subspace of a *PN*-dimensional space, i.e. 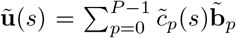 with 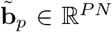. In particular, we set 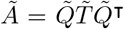 with

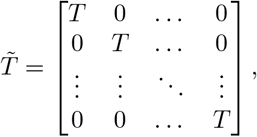

and with 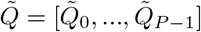 with 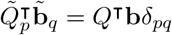 and 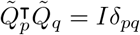. Then we have:

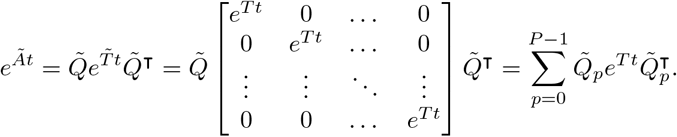

We can then compute the signal-to-noise ratio for any stimulus pair

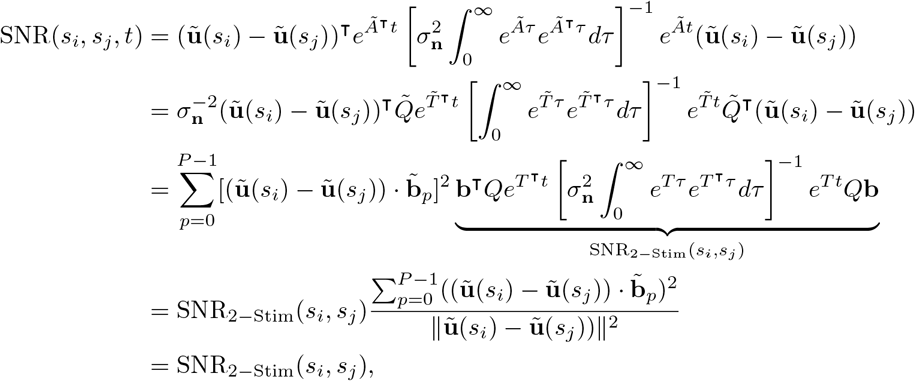

where SNR_2*−*Stim_(*s*_*i*_, *s*_*j*_) is the SNR of the original two-stimulus network *A* for discrimination of a pair of inputs with 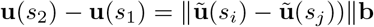.

Thus, by aligning an orthogonal subspace to each stimulus dimension 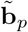, networks optimised for discrimination of a single stimulus pair can be naturally composed to form networks that discriminate arbitrarily many stimuli drawn from a *P*-dimensional space, with the number of neurons scaling only linearly with *P*.

### G Targeted Dimensionality Reduction

Given a set of measured responses **x**^(*k*)^(*t*) of *N* neurons, over *K* trials with stimuli stim(*k*) = *θ*^(*k*)^, and *T* time points per trial, we fit the model:

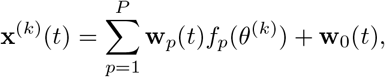

where **w**_*p*_(*t*) are neuron- and time-dependent factors and *f*_*p*_(*θ*^(*k*)^) are stimulus-dependent factors. This equation can be written in matrix form as:

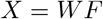

where

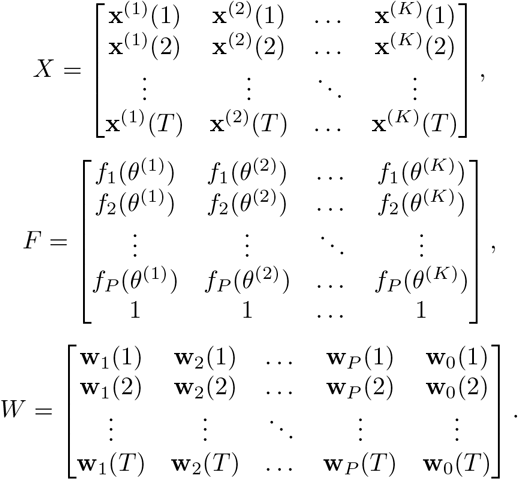

Thus, given an estimate of the set of *P* vectors 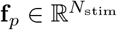, a least squares estimate of the factors **w**_*p*_(*t*) can be obtained by solving the above linear equation via the Moore-Penrose pseudoinverse *W* = *XF*^+^.

To solve for *f*_*p*_(*θ*) given estimates of **w**_*p*_(*t*), note that:

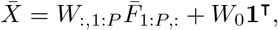

where 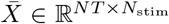 contains the PSTHs 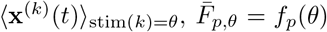 and 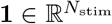 is a vector of ones (with 1 ≤ *θ* ≤ *N*_stim_ the stimulus labels). Thus, we obtain 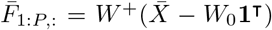. To fit the TDR model to data, we iterated between these two updates until convergence, initialising with *f*_*p*_(*θ*) ∼ *N*(0, 1) (for 1 ≤ *p* ≤ *P* and 1 ≤ *θ* ≤ *N*_stim_).

The TDR model has a number of invariances that we exploited to convert the model into a standard form. First, the stimulus factors can be written as 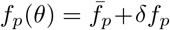 where 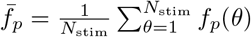 is the mean over stimuli. We removed the mean component for all *p* by setting 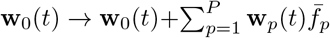.Second, there exists an invariance 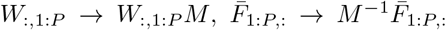 for arbitrary invertible *M* ∈ *ℝ*^*P ×P*^. We selected a unique basis by taking the singular value decomposition 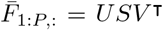 and redefining 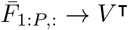 and *W*_:,1:*P*_ → *W*_:,1:*P*_ *US* (note that this was called the Principal Component Basis by Aoi et al. (2020)). An advantage of this choice of basis is that the vectors **f**_*p*_ are orthonormal, and the stimulus-independent component **f**_0_ = **1** is also orthogonal (since all other vectors have zero mean, and the dot product of a constant vector with a zero mean vector is zero). Finally, for the circular stimulus set in the working memory task, *F*_1:*P*,:_ had two leading principal components of roughly equal magnitude (the two basis vectors that spanned the circle), so that the Principal Component Basis was approximately invariant to a further rotation in this two-dimensional subspace. To ensure that all models (with *P* = 2) were plotted and compared in a comparable basis, we therefore performed an Orthogonal Procrustes transformation to obtain a 2 × 2 orthogonal matrix 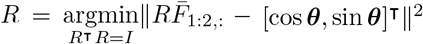 and set 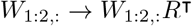 and *W*_1:2,:_ → *W*_1:2,:_*R*^⊺^.

## Supplementary Figures

**Figure S1:**
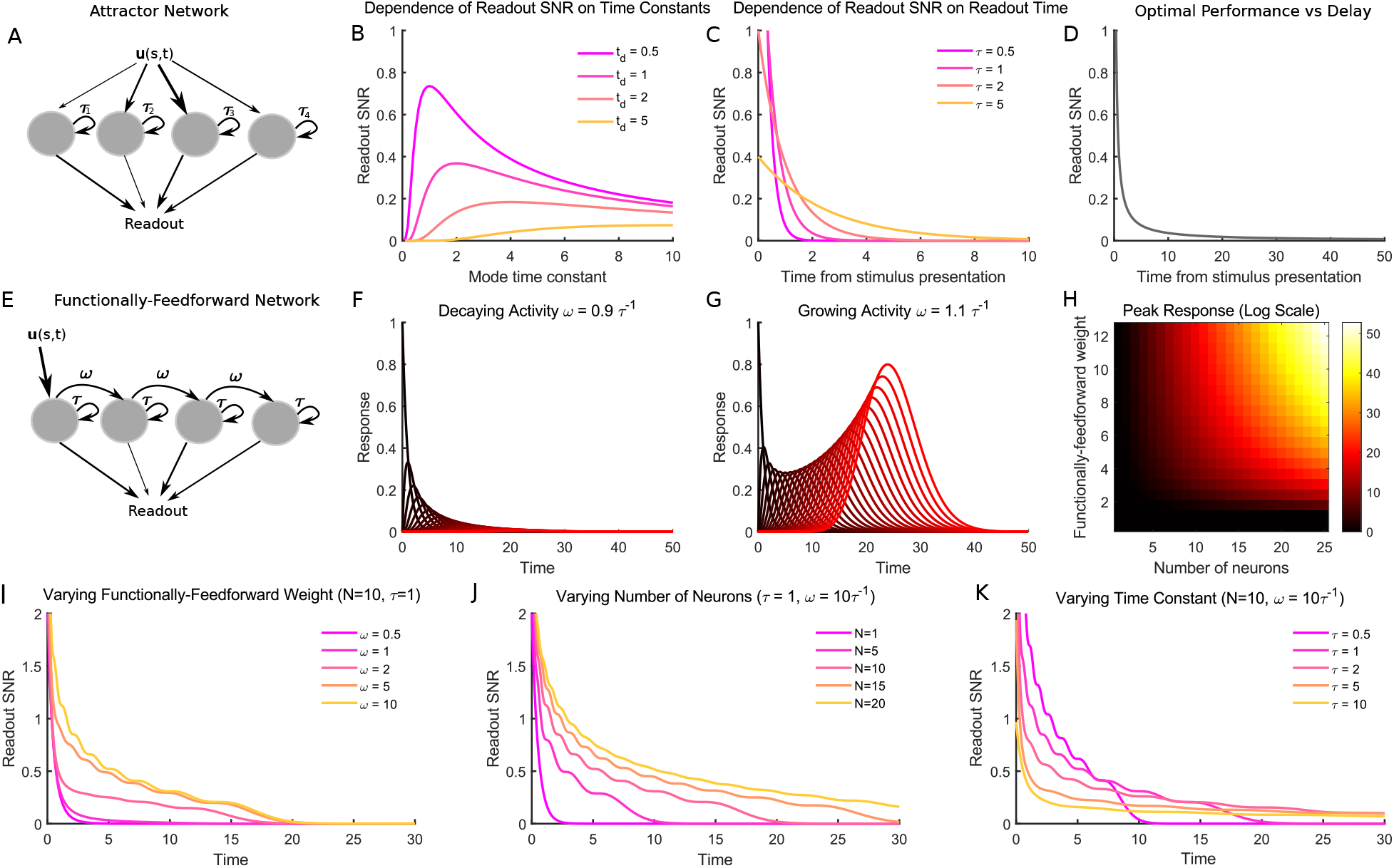
Behaviour of attractor (normal) and functionally-feedforward (non-normal) networks on the WM task with delta pulse stimulus input and stationary state covariance. A: Normal networks can be characterised by a set of independent eigenmodes. Each eigenmode applies an independent filter 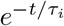 to the projection of network input onto its left eigenvector 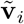. A downstream readout could in principle combine information from multiple such eigenmodes to obtain information about different projections of the recent history of sensory input over different timescales. B: A single eigenmode can be optimised for readout at a specific delay time *t*_*d*_ by setting its time constant as *τ*_*k*_ = 2*t*_*d*_. C: Readout performance along a single eigenmode decays exponentially after stimulus presentation. D: Even when optimised for a given delay, the readout performance decays as (*et*_*d*_)^*−*1^ for attractor networks. E: Functionally-feedforward networks are better characterised by a Schur decomposition, in which each mode has a decay time constant *τ* and a feedforward weight onto the next mode *ω* (also called a homogeneous delay line). F, G: Activity propagates along the delay line, and depending on the value of *ωτ*, can either decay (*ωτ <* 1, F) or grow (*ωτ >* 1, G) exponentially along the delay line. H: Despite being asymptotically stable when *τ >* 0, activity can grow to arbitrarily large values as *ω* and *N* are increased. I-K: The performance of the functionally-feedforward network grows as *ω* and *N* are increased and *τ* is decreased, with SNR → ∞ when *N, ω* → *∞* and *τ* → 0. Thus, the delay line can substantially outperform normal networks (A-D), but these improvements require an exponential increase in the firing rate of neurons to achieve (H).

**Figure S2:**
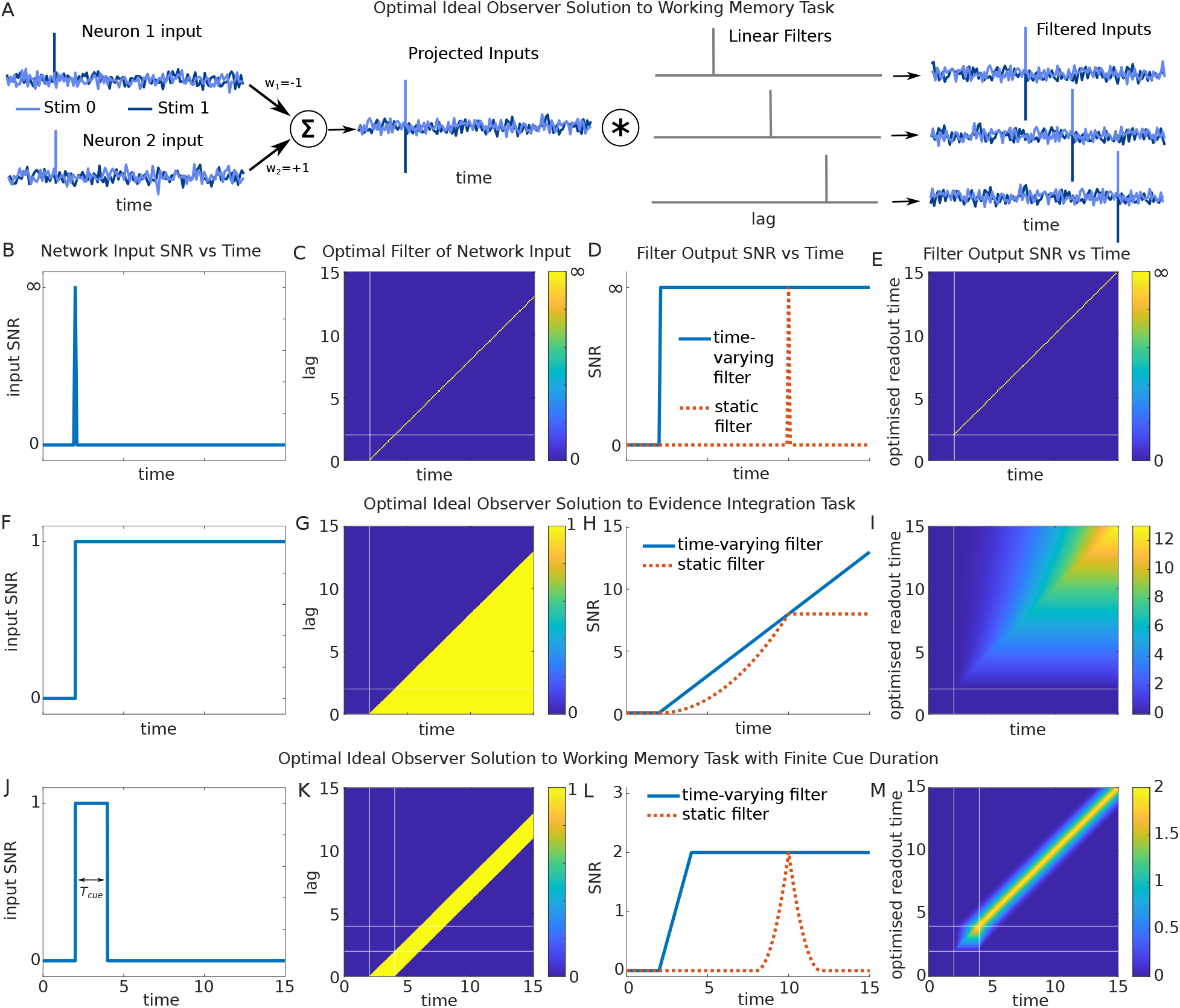
Bayesian deal observer solutions to WM and perceptual decision-making tasks. A: Stimuli can be optimally discriminated based on network input by first projecting these inputs onto their linear discriminant, then convolving them with a delay filter matched to the readout time *t*_*d*_ and applying a threshold operation (not shown). B: Signal-to-noise ratio of network input vs time for the WM task with delta pulse stimulus input. C: A linear time-varying linear filter *f*(*t*_*d*_, *τ*) = *δ*(*t*_*d*_ − *t*_*s*_ − *τ*) achieves the optimal response SNR. D: The SNR of filter output is infinite at the optimised readout time *t*_*d*_. The optimal time-varying filter achieves perfect performance at all times *t > t*_*s*_, while a static filter can achieve optimal performance only at the optimised delay *t*_*d*_. E: The SNR of filter output vs time for static filters optimised for each readout time. F: In a perceptual decision-making task in which the stimulus is presented continuously, the SNR of network input is zero before stimulus onset and constant after stimulus onset, requiring temporal integration to achieve a high output SNR. G: The optimal time-varying filter for this task integrates the history of network input for all times *t > t*_*s*_, but assigns zero weight to inputs at times *t < t*_*s*_. H: Response SNR for the optimal time-varying filter grows linearly after stimulus onset, while an optimal static filter has bounded performance. J-M: When the stimulus appears for a fixed interval of time *T*_cue_, the optimal filter integrates network input during the stimulus period (with an integration window *T*_cue_) and then delays this input for a period *t*_*d*_ − *t*_*s*_ − *T*_cue_.

**Figure S3:**
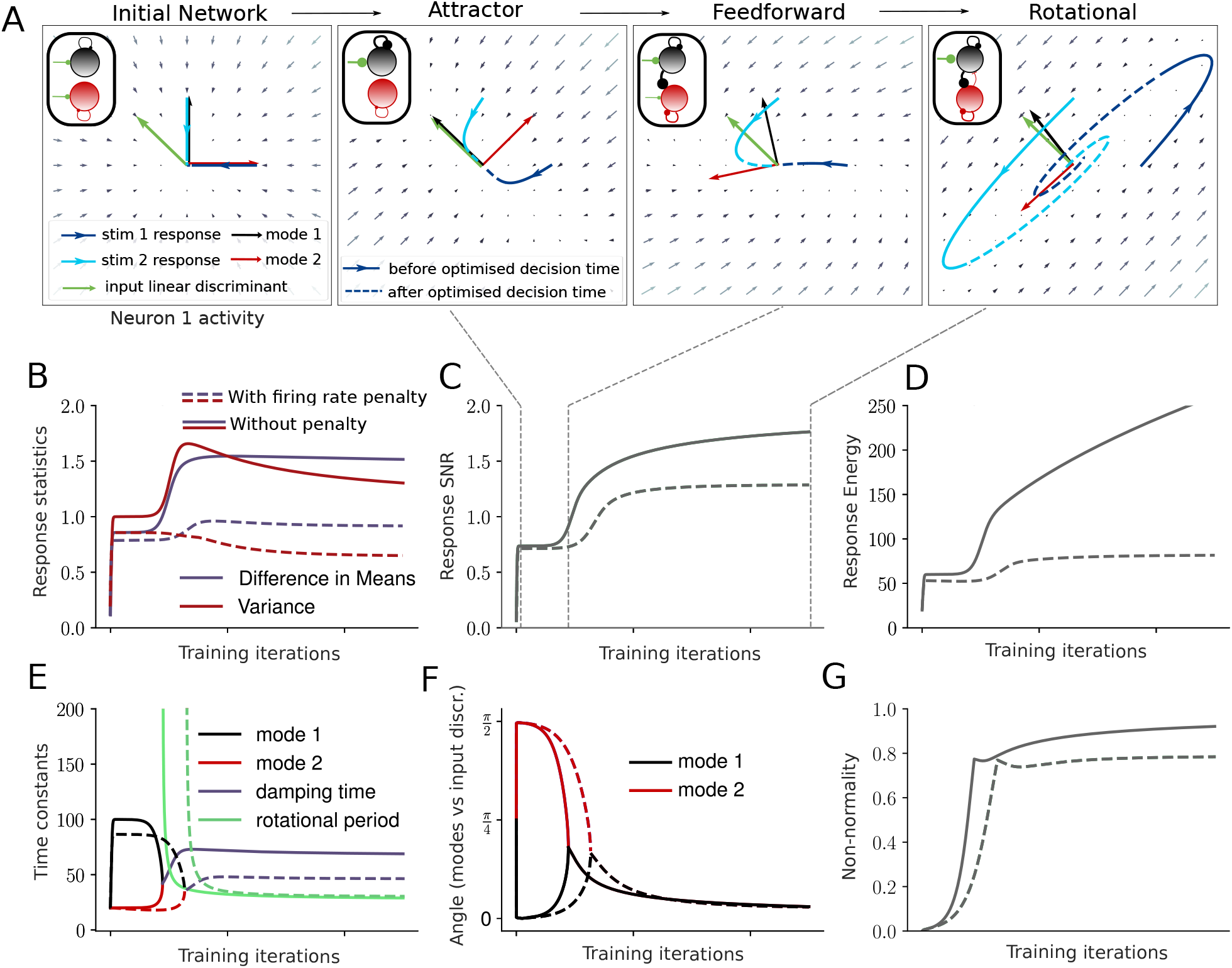
Influence of energetic penalty on learned dynamics. A: As in Figure 2C, but showing the linear discriminant and two dynamical modes (Schur modes) at each stage of optimisation (without energy penalty). B: The mean separation and variance of responses along the readout vector at the decision time, at each iteration of the optimisation process for networks with and without a penalty on energetic cost. C: The SNR of responses for the two networks. D: The total energy of the network response (time-integrated squared norm of firing rate vector). E: The time constants of the two dynamical modes. Before the transition to rotational dynamics, the time constants are determined by the (real) eigenvalues. After the transition, they are given by the real and imaginary parts of a complex-conjugate pair of eigenvalues. F: The angle between each input mode (left eigenvector) and the input linear discriminant. After the transition to rotational dynamics the input Schur mode is shown. G: Non-normality of the dynamics matrix *A* (normalised between 0 and 1, see Methods).

**Figure S4:**
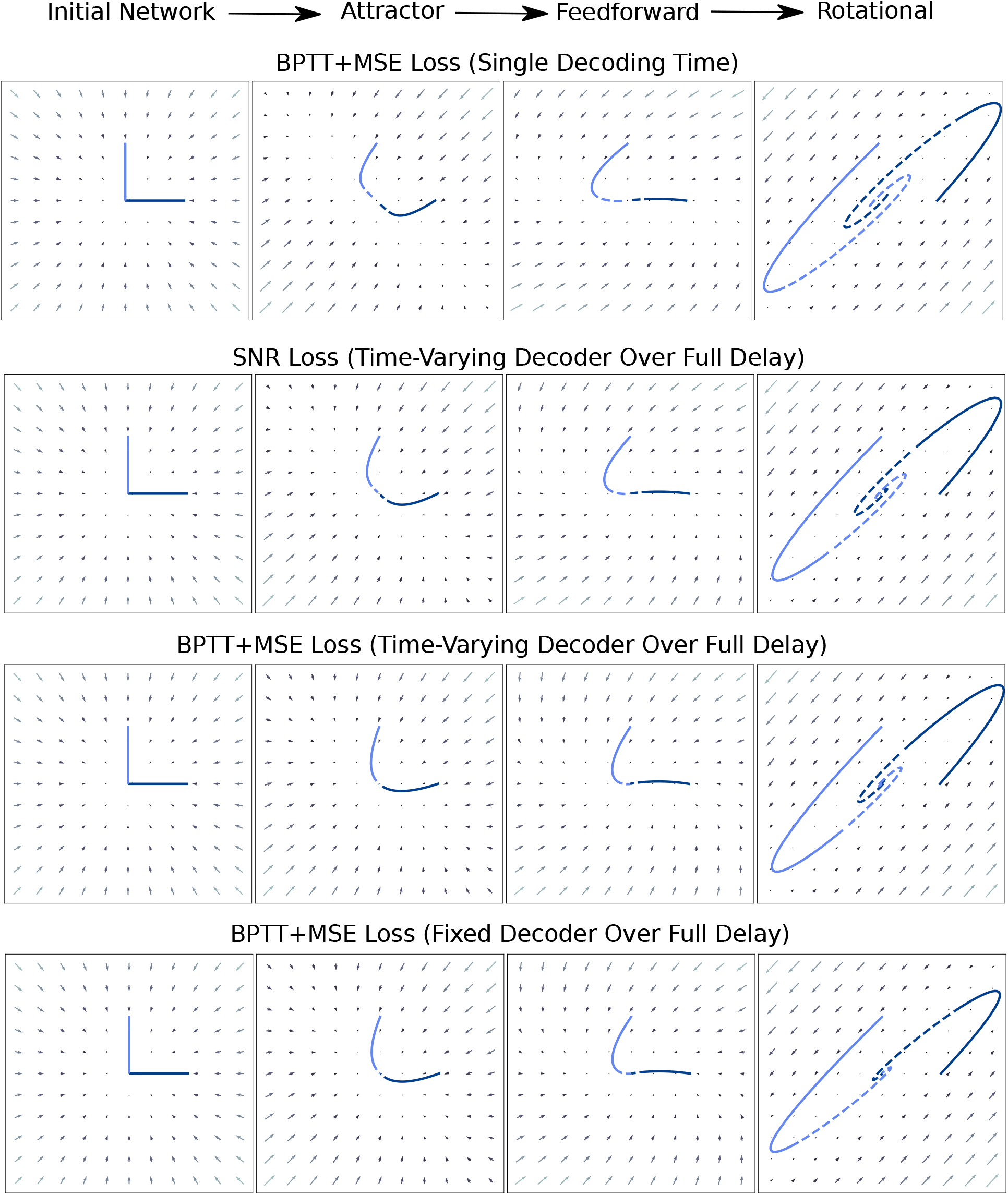
Comparison of networks optimised using 1) SNR loss vs backpropagation through time (BPTT) with mean squared error (MSE) loss, 2) loss evaluated at a single time point vs all time points over the delay, and 3) a fixed decoder vs a time-varying decoder. Top row: Network optimised using BPTT with MSE loss at a single time point *t*_*d*_ = 50. Second row: Optimisation with the SNR loss, weighted equally at all delays 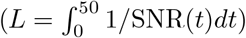. Third row: optimisation using BPTT+MSE with a separate linear decoder fit to each time point *t*_*d*_ ∈ [0, 50]. Bottom row: Optimisation using BPTT+MSE with a single decoder fit to minimise MSE across all time points *t*_*d*_ ∈ [0, 50].

**Figure S5:**
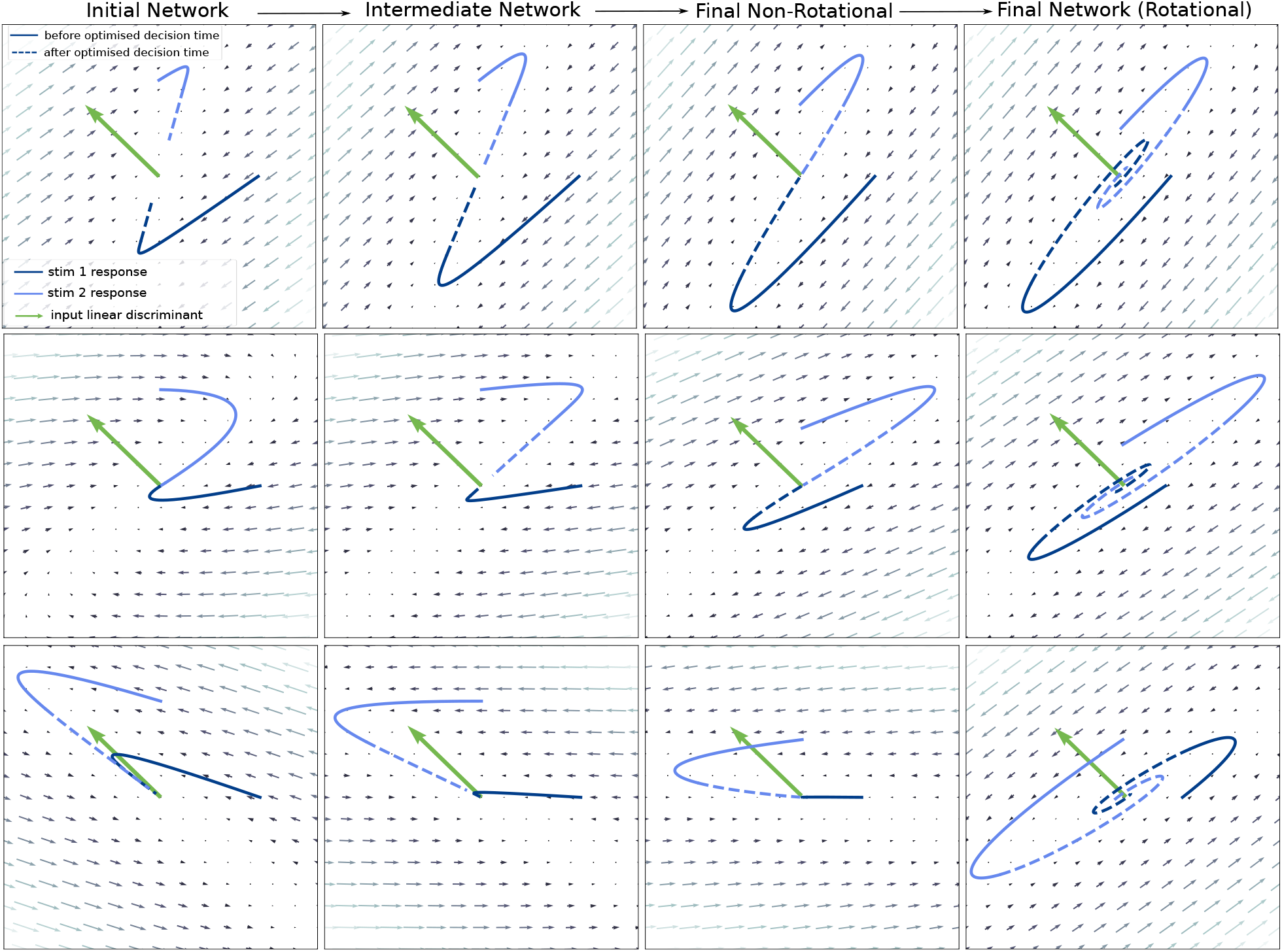
Rotational solutions are insensitive to initial network weights. Each row shows four phases of the optimisation of a network with different random weight initialisation.

**Figure S6:**
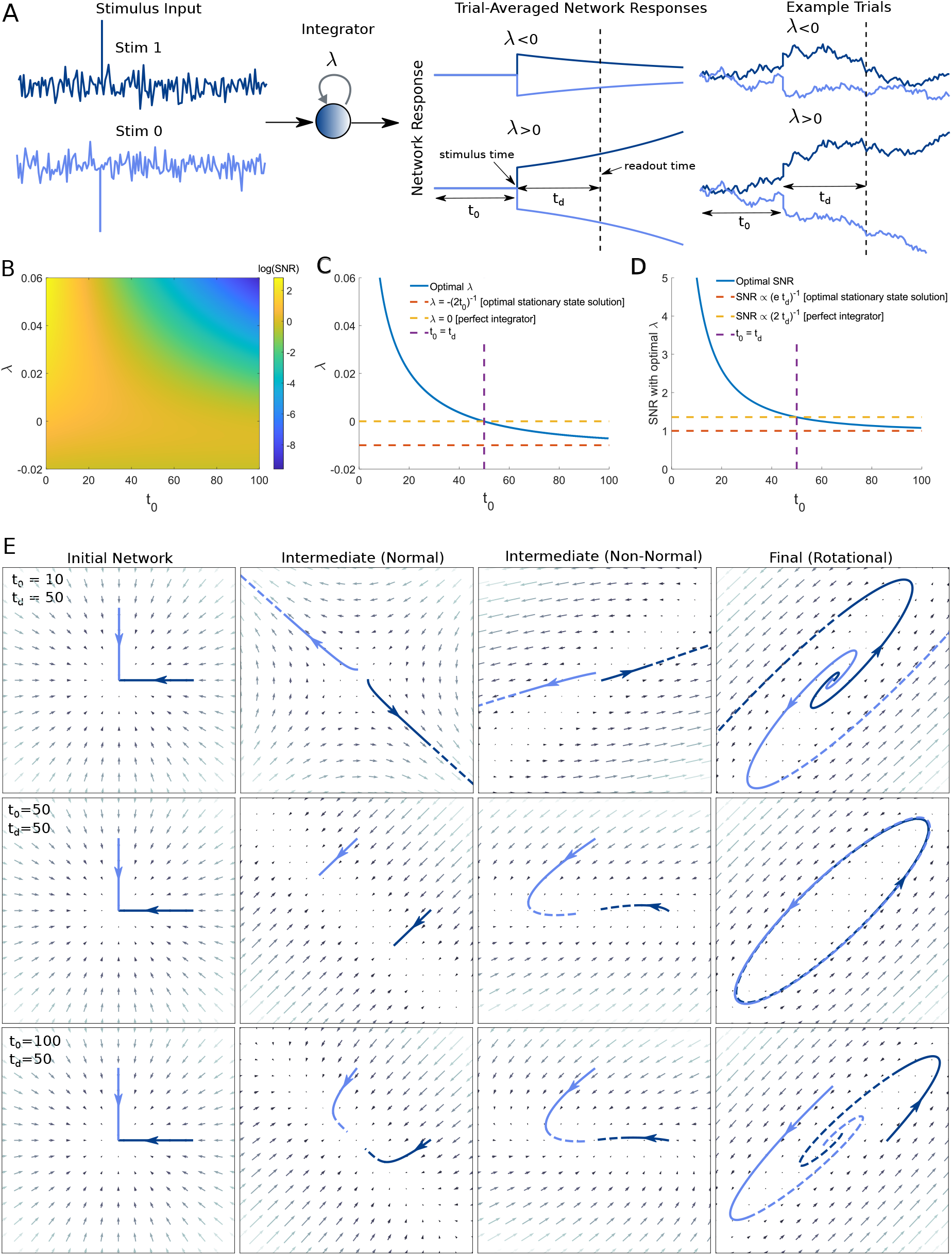
Effect of initial state covariance on optimal dynamics. A: We derived analytical solutions for a one-dimensional integrator with recurrent weight *λ* receiving inputs from one of two stimuli given by *u*(*s*_0_, *t*) = −*δ*(*t* − *t*_*s*_) + *n*(*t*) and *u*(*s*_1_, *t*) = *δ*(*t* − *t*_*s*_) + *n*(*t*). The integrator is inititialised at a fixed (zero variance) initial condition at a time *t*_*s*_ −*t*_0_ and the readout occurs at time *t*_*s*_ +*t*_*d*_, where *t*_*s*_ is the stimulus time. Thus, noise accumulates for a time window *t*_0_ before stimulus onset. B: The response SNR at the readout time depends on the values of *t*_0_ and *λ*. C, D: The optimal value of *λ* is positive for *t*_0_ *< t*_*d*_ and negative for *t*_0_ *> t*_*d*_, reflecting a transition from leaky integrator to unstable amplifier as the pre-stimulus noise integration time window *t*_0_ is decreased. E: Numerically optimised networks exhibit rotational dynamics, but the damping rate exhibits the predicted transition from stable and

**Figure S7:**
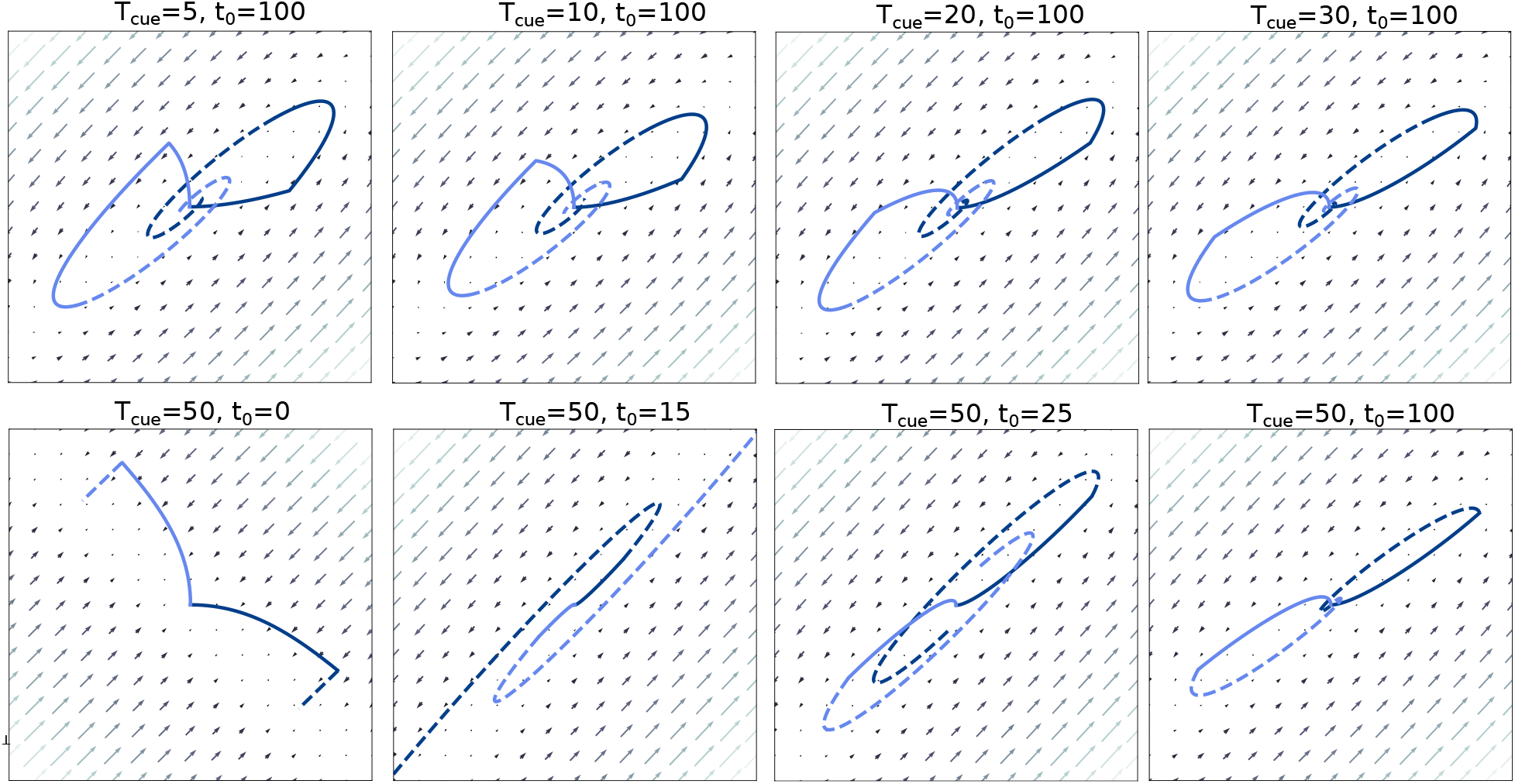
Effect of length of stimulus period and initial state covariance on optimal dynamics. Top row shows effect of changing the length of the stimulus period *T*_cue_. The readout time was *t*_*d*_ = 50 and networks were driven by noise for a period *t*_0_ = 100 before stimulus onset (at *t*_*s*_ = 0). All optimised networks had complex eigenvalues, and the differences in trajectories were primarily due different input durations *T*_cue_ rather than changes in flow fields. Bottom row shows effect of varying the duration of pre-stimulus noise accumulation *t*_0_ for networks with the stimulus presented continuously from *t* = 0 to *t* = *t*_*d*_ = 50. The network optimised with *t*_0_ = 0 learned a classic line attractor solution to the task, which is known to be optimal in this setting (Gold and Shadlen, 2007). In contrast, for *t*_0_ *>* 0 a line attractor would generate substantial pre-stimulus variability, and so is no longer optimal. In this case, networks must trade off integration during stimulus presentation against avoidance of integration of pre-stimulus noise (Supplementary Figure S2F-I). All networks optimised with *t*_0_ *>* 0 had complex eigenvalues, suggesting that rotational dynamics are optimal for evidence integration tasks in which pre-stimulus noise influences task performance. Note that, although the trajectories during the stimulus presentation (solid lines) use the linear part of the curved rotational trajectories, noise inputs before stimulus onset would have been integrated for a longer period of time and therefore would be rotated through the curved part of the trajectories (dashed lines), thereby reducing their influence on the decoder.

**Figure S8:**
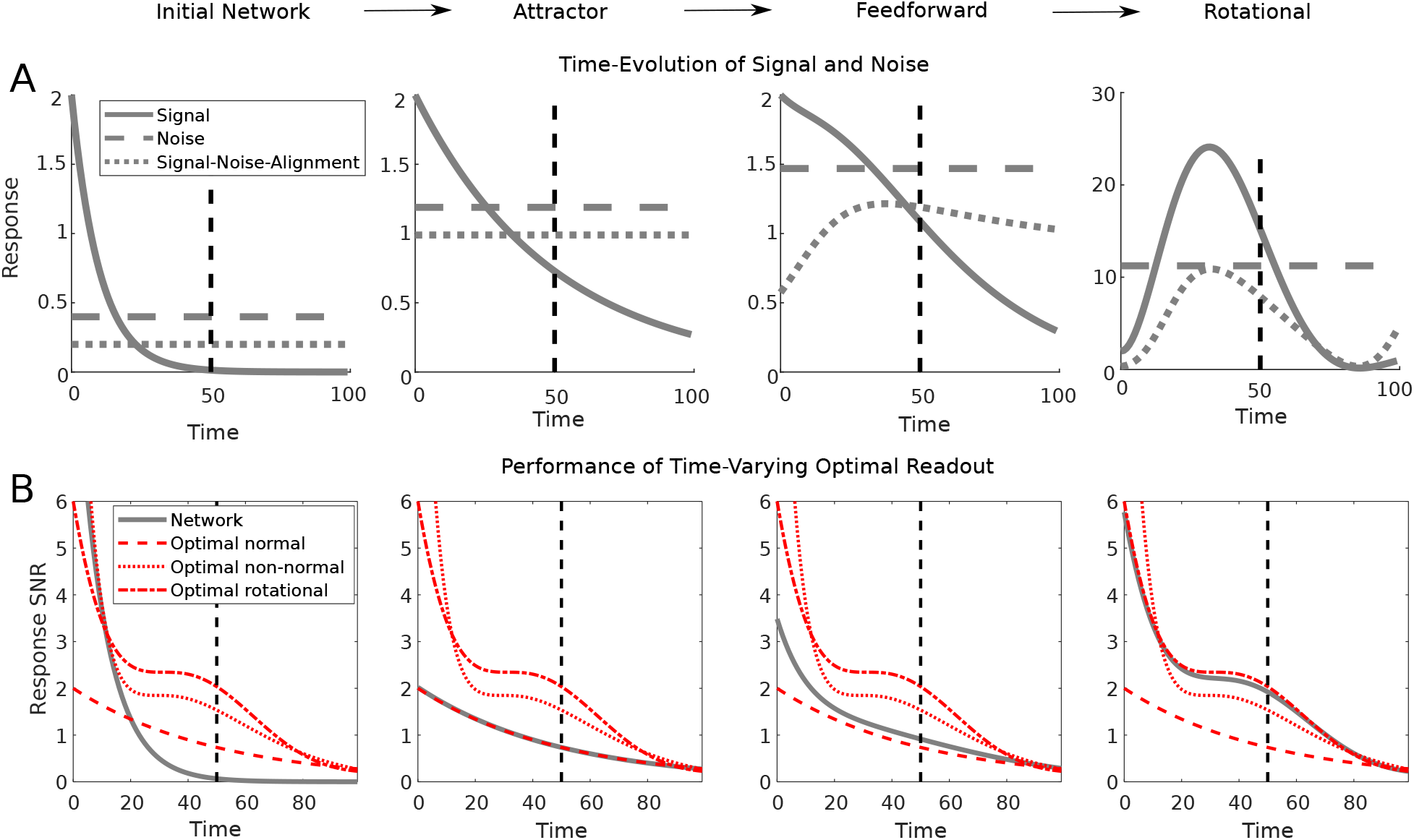
A: The signal (squared norm of vector separating two stimulus-evoked mean responses), noise (total variance of responses to either stimulus) and signal-noise-alignment (see Methods) for each network as a function of time during the delay. Vertical dashed black line shows optimised decision time *t*_*d*_. Note that networks were initialised at stationary state, so that noise statistics do not vary during the delay. B: The performance of the optimal readout at each time following stimulus onset (grey line). Red lines show the analytically computed optimal normal and two-dimensional non-normal or rotational networks.

**Figure S9:**
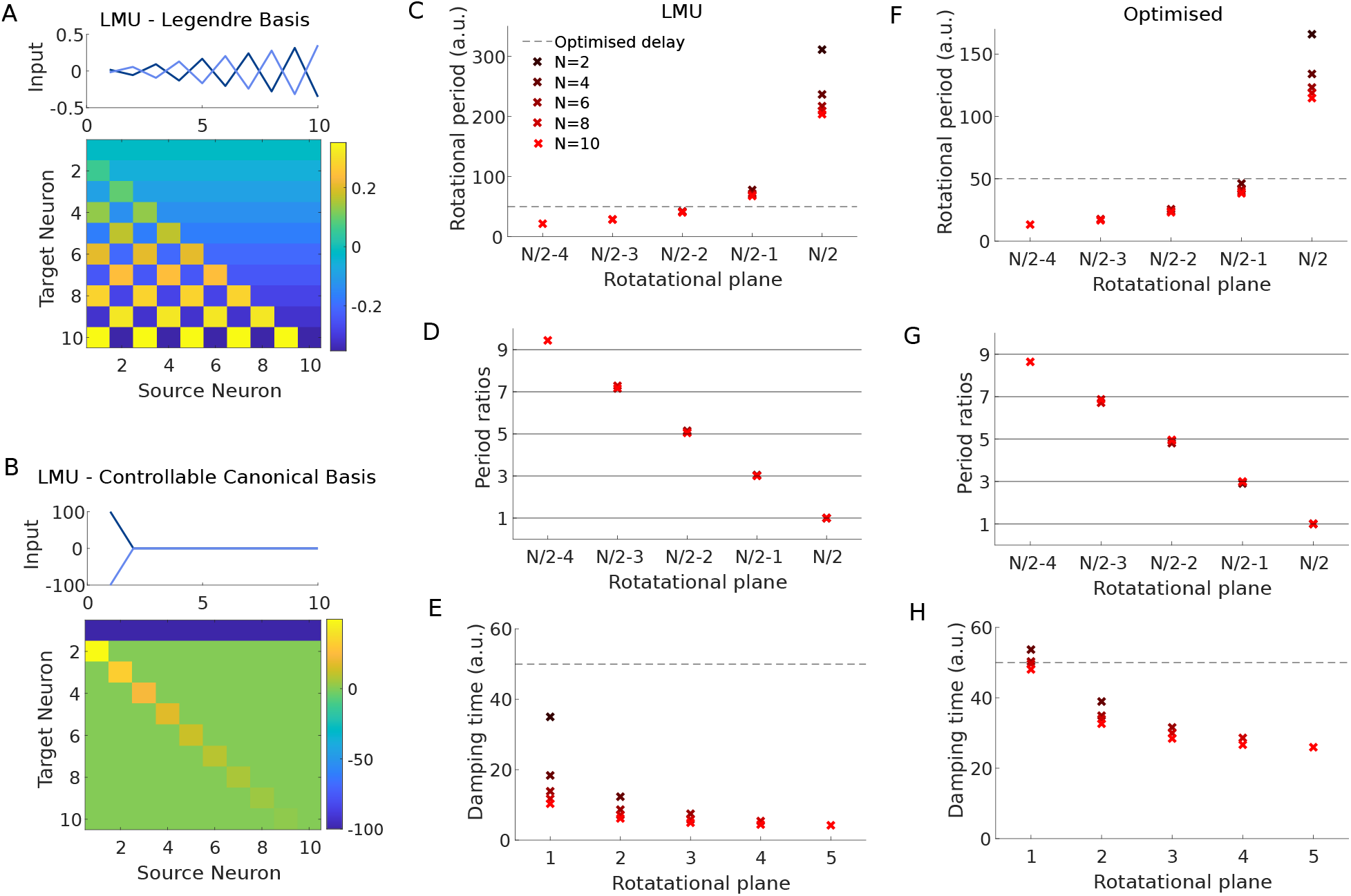
Rotational structure in optimised networks vs the Legendre Memory Unit (LMU). A: Representation of the LMU in the Legendre basis. B: Equivalent representation of the LMU in the Controllable Canonical Basis. C: The rotational periods (given by imaginary parts of eigenvalues) of the LMU. Note that the eigenvalues are the same in the two bases in A and B. D: The ratio of each rotational period with that of the output rotational plane. E: The damping times (given by real parts of eigenvalues) of the LMU. F-H: As in C-E, but for the numerically optimised networks.

**Figure S10:**
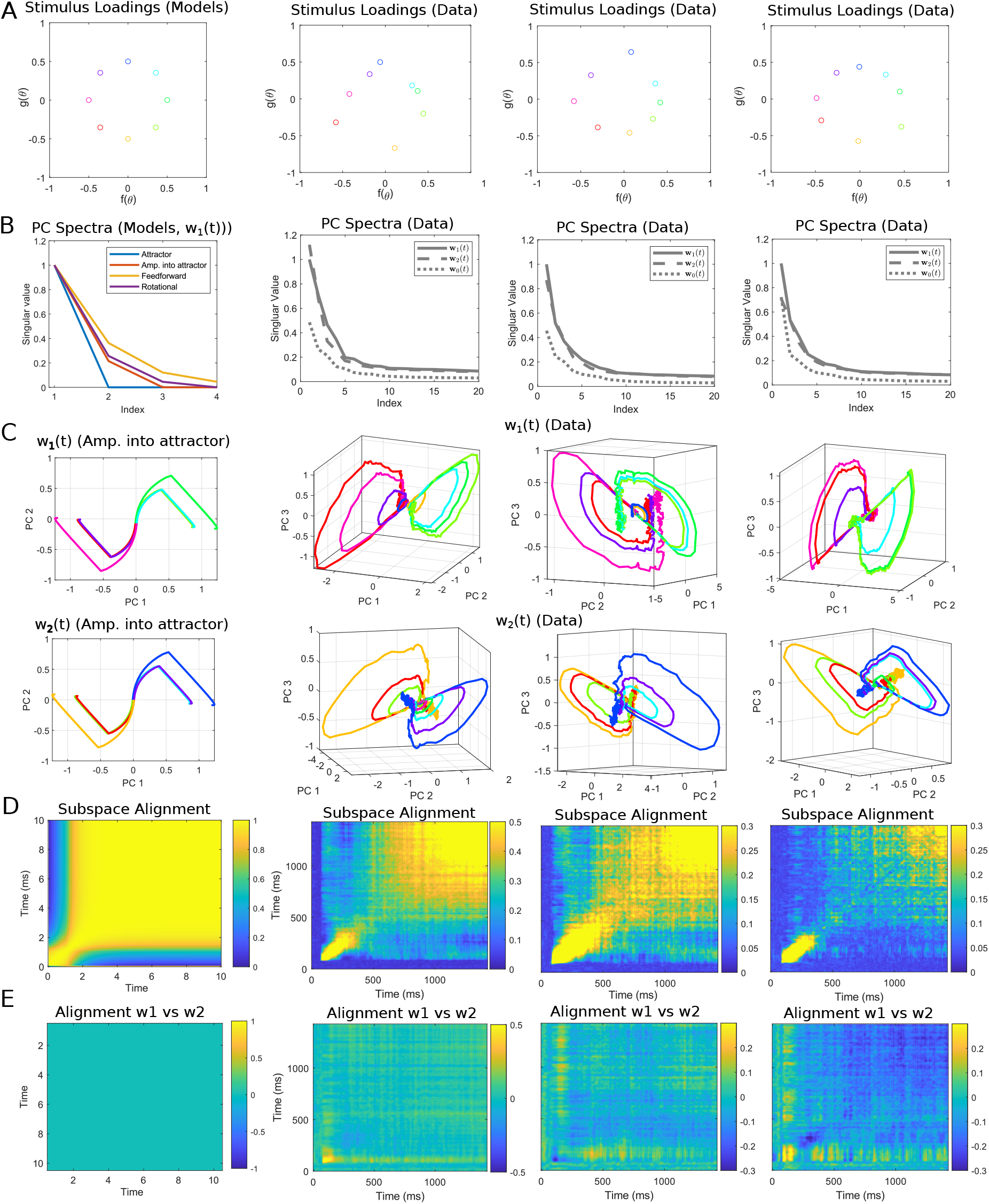
A: Stimulus loadings *f*(*θ*), *g*(*θ*) in TDR models fit to data simulated from each WM model (left panel, all models yield identical results), and three experimental recording sessions (right three panels). B: Principal component spectrum of **w**_*p*_(*t*) matrices of TDR models fit to simulated data (left panel) and three experimental sessions (right three panels). For the simulated data, only **w**_1_(*t*) is shown, but **w**_2_(*t*) has an identical spectrum and **w**_0_(*t*) = 0. For the experimental recordings, all three matrices are shown, with singular values normalised by the maximum singular value of **w**_1_(*t*). C: Responses to each stimulus along the top principal components of **w**_1_(*t*) and **w**_2_(*t*) for the amplification into attractor model (left) and three recording sessions. Other models are shown in Figure 5. D: The alignment of the plane spanned by **w**_1_(*t*), **w**_2_(*t*) and the plane spanned by **w**_1_(*t*^*′*^), **w**_2_(*t*^*′*^) (left: amplification into attractor model, right: data). E: Alignment of **w**_1_(*t*) and **w**_1_(*t*^*′*^) (left: amplification into attractor model, right: data).

## Footnotes

1 Sylvester equations have a unique solution if *P* and *−Q* do not share a common eigenvalue. Here, however, *P* = *−Q*, violating this condition.

2 We choose a lower-triangular rather than upper-triangular decomposition for this analysis as it leads to simpler indices in the following derivations.

